# A widespread hydrogenase drives fermentative growth of gut bacteria in healthy people

**DOI:** 10.1101/2024.08.15.608110

**Authors:** Caitlin Welsh, Princess R. Cabotaje, Vanessa R. Marcelino, Thomas D. Watts, Duncan J. Kountz, Jodee A. Gould, Nhu Quynh Doan, James P. Lingford, Jessica Solari, Gemma L. D’Adamo, Ping Huang, Natasha Bong, Emily L. Gulliver, Remy B. Young, Kaija Walter, Patricia G. Wolf, Jason M. Ridlon, H. Rex Gaskins, Edward M. Giles, Dena Lyras, Rachael Lappan, Gustav Berggren, Samuel C. Forster, Chris Greening

## Abstract

Molecular hydrogen (H_2_) is among the most central, but least understood, metabolites in the human gastrointestinal tract (gut). H_2_ gas is produced in large quantities during bacterial fermentation and consumed as an energy source by bacteria and archaea. Disruption of H_2_ cycling is linked to gastrointestinal disorders, infections, and cancers, with H_2_ used as an indicator of gut dysfunction through breath tests. Despite this, the microorganisms, pathways, and enzymes mediating H_2_ production remain unresolved. Here we show that a previously uncharacterised enzyme, the group B [FeFe]-hydrogenase, drives most fermentative H_2_ production in the human gut. Analysis of stool, biopsy, and isolate (meta)genomes and (meta)transcriptomes show this hydrogenase is encoded by most gut bacteria and is highly expressed. Through analysis of 19 taxonomically diverse gut isolates, the group B [FeFe]-hydrogenase produces large amounts of H_2_ gas and supports fermentative growth of both Bacteroidetes and Firmicutes. *Bacteroides* particularly dominate H_2_ production. Biochemical and spectroscopic characterisation shows purified group B [FeFe]-hydrogenases are catalytically active and bind a di-iron active site. These hydrogenases are highly enriched in the guts of healthy individuals, but significantly depleted in favour of other fermentative hydrogenases in Crohn’s disease. Furthermore, we show that metabolically flexible respiratory bacteria are the most abundant H_2_ oxidizers in the gut, not sulfate reducers, methanogens, and acetogens as previously thought. This combination of enzymatic, cellular, and ecosystem-level analysis provides the first detailed understanding of H_2_ cycling in the human gut and reveals new links between microbiota function and gastrointestinal health.

## Introduction

Molecular hydrogen (H_2_) is a central intermediate in gastrointestinal digestive processes. Most bacteria within the gut hydrolyse and ferment dietary carbohydrates to absorbable short-chain fatty acids^1–3^ and large quantities of H_2_ gas^4,5^. H_2_ accumulates to high micromolar levels in the gut, where it is primarily consumed by other microbes for energy conservation and carbon fixation^6,7^, though some is also expelled as flatus or exhaled^8–10^. Classically, three groups of gut microbes are thought to consume H_2_, namely acetogenic bacteria, methanogenic archaea, and sulfate-reducing bacteria^3,11–14^. H_2_ consumption by gut microbes lowers H_2_ partial pressures, thereby ensuring fermentation remains thermodynamically favourable^3,15–19^. In turn, many H_2_-producing and H_2_- consuming microbes form mutualistic relationships by conducting interspecies H_2_ transfer depending on physical association^15,20^. In addition to supporting digestion, gastrointestinal H_2_ cycling modulates levels of important metabolites in the gut, including butyrate^21^, hydrogen sulfide^22^, bile acids^2^, and host steroids^23^, with diverse effects on processes such as digestion, inflammation, and carcinogenesis. It is also proposed that microbiota-derived or therapeutically-supplied H_2_ may directly benefit human cells as an antioxidant^24,25^. Disruption of the balance between H_2_-producing and H_2_-consuming bacteria has been linked with a range of gut and wider disorders^19,26^; most notably, gas buildup contributes to the symptoms of irritable bowel syndrome (IBS) and hydrogen breath tests are frequently, if controversially, used to detect disorders such as carbohydrate malabsorption^27–29^. Numerous pathogens also exploit microbiota-derived H_2_ during invasion, including *Helicobacter pylori* and *Salmonella*^30–34^, or rapidly produce it in the case of pathogenic Clostridia and protists^30,35,36^.

Despite the central importance of H_2_ cycling in human health and disease, surprisingly little is known about which microbes and enzymes mediate this process. Both the production and consumption of H_2_ are catalysed by hydrogenases, which fall into three major groups dependent on the metal content of their active site, the [FeFe]-, [NiFe]-, and [Fe]-hydrogenases, and multiple subgroups^37,38^. It’s been classically thought that most H_2_ production in the gut is mediated by fermentative bacteria, primarily the class Clostridia, that couple reoxidation of ferredoxin (e.g. reduced during acetate fermentation by the pyruvate-ferredoxin oxidoreductase reaction) to the evolution of H_2_. Some Clostridia use group A1 [FeFe]-hydrogenases, an extensively structurally and mechanistically characterised lineage of enzymes, to rapidly produce H_2_^39,40^. Some H_2_ may also be produced by formate hydrogenlyase complexes (containing a group 4a [NiFe]-hydrogenase) that disproportionate formate during fermentative survival of Enterobacteriaceae^41,42^. Yet two recent findings suggest that other fermenters are also active in the human gut. Our 2016 survey showed a distantly related enzyme (28% amino acid identity) called the group B [FeFe]-hydrogenase is widespread in diverse gut isolates and abundant in gut metagenomes ^19,43^. Two recent biochemical studies suggest these enzymes are active and biased towards H_2_ production^44,45^, though their physiological activity and role has yet to be confirmed in any organism. In parallel, trimeric electron-confurcating hydrogenases (group A3 [FeFe]-hydrogenases) have been discovered that couple oxidation of NAD(P)H and reduced ferredoxin to the evolution of H_2_ ^46–49^. We have demonstrated that group A3 [FeFe]- hydrogenases are primarily responsible for H_2_ production in ruminants^17,50,51^, though it is unclear if these principles also extend to humans. Similarly, it is unclear whether the paradigms regarding H_2_ consumption are accurate, given the three classical groups of H_2_ oxidizers (hydrogenotrophs) are generally in low abundance in the human gut. Indeed, only approximately half of people produce methane gas^52,53^ and it is becoming increasingly apparent that most hydrogen sulfide is derived from organosulfur compounds rather than sulfate reduction^54–56^. Respiratory hydrogenotrophs that use electron acceptors such as fumarate, nitrate, sulfoxides, and inflammation-derived oxygen may also be active but overlooked members of gut microbiota^19,31^.

Here we gained the first detailed understanding of the enzymes and microbes responsible for hydrogen cycling in the human gastrointestinal tract. To do so, we holistically profiled the abundance, expression, and distribution of hydrogenases using metagenomes and metatranscriptomes, including original biopsy samples, in both healthy individuals and those with gastrointestinal disorders. We then performed in-depth analysis of 19 bacterial isolates and four heterologously produced enzymes to confirm the activity and roles of these enzymes. We reveal that group B [FeFe]- hydrogenase drives most H_2_ production in the human gut, highlight the overlooked role of *Bacteroides* as major H_2_-producing fermenters, and show that hydrogenases are differentially abundant between healthy people and those with chronic disease phenotypes such as Crohn’s disease.

## Results and Discussion

### Group B [FeFe]-hydrogenases are the most widespread and expressed gut hydrogenases

We initially investigated the distribution of hydrogenase genes in the human gut by analysing 300 human stool metagenomes^57^ **(Table S1)**. Hydrogenase genes are extremely abundant, occurring on average at 1.44 ± 0.58 copies per genome (cpg), based on normalisation to single-copy bacterial and archaeal marker genes **(Fig. 1a)**. By far the most abundant are the functionally uncharacterised group B [FeFe]-hydrogenase (0.75 ± 0.25 cpg), hypothesized but unproven to mediate fermentative H_2_ production^19^ **(Fig. 1b)**. Genes encoding these enzymes are much more abundant than the monomeric ferredoxin-dependent group A1 [FeFe]-hydrogenases (0.10 ± 0.09 cpg), previously thought to account for most fermentative hydrogen production in gastrointestinal tracts^6,58,59^, and trimeric electron-confurcating group A3 [FeFe]-hydrogenases (0.19 ± 0.11 cpg) that dominate H_2_ production in ruminants^17,50^. Other enzymes also potentially play minor roles in H_2_ production in the human gut, including formate hydrogenlyases (group 4a [NiFe]-hydrogenases; 0.02 ± 0.07 cpg) and possibly ferredoxin-dependent energy-converting hydrogenases (group 4e [NiFe]-hydrogenases; 0.06 ± 0.04 cpg) **(Table S1; Fig. 1a & 1b)**. Consistently, analyses of 78 paired metatranscriptomes confirm that these hydrogenases are highly expressed (RNA / DNA ratios between 1.76 to 6.90 depending on subgroup) **(Table S1; Fig. 1a)**. Transcripts for the group B [FeFe]-hydrogenase are the most numerous (95 ± 86 reads per kilobase per million mapped reads, RPKM; RNA / DNA ratio = 2.0) and 3.3-, 4.7-, and 26-fold higher than the group A3, A1, and 4a enzymes typically thought to be responsible for gastrointestinal H_2_ production **(Fig. 1a)**. Given [FeFe]-hydrogenases are usually highly active enzymes^39,40^, these expression levels are likely to enable rapid H_2_ production in the gut. Nitrogenases, which produce H_2_ during their reaction cycle^60^, were also widely encoded but minimally expressed by gut bacteria **(Fig. 1a)**. Altogether, group B [FeFe]-hydrogenases potentially drive most H_2_ production in the gut, though operate alongside other enzymes.

**Figure 1.**
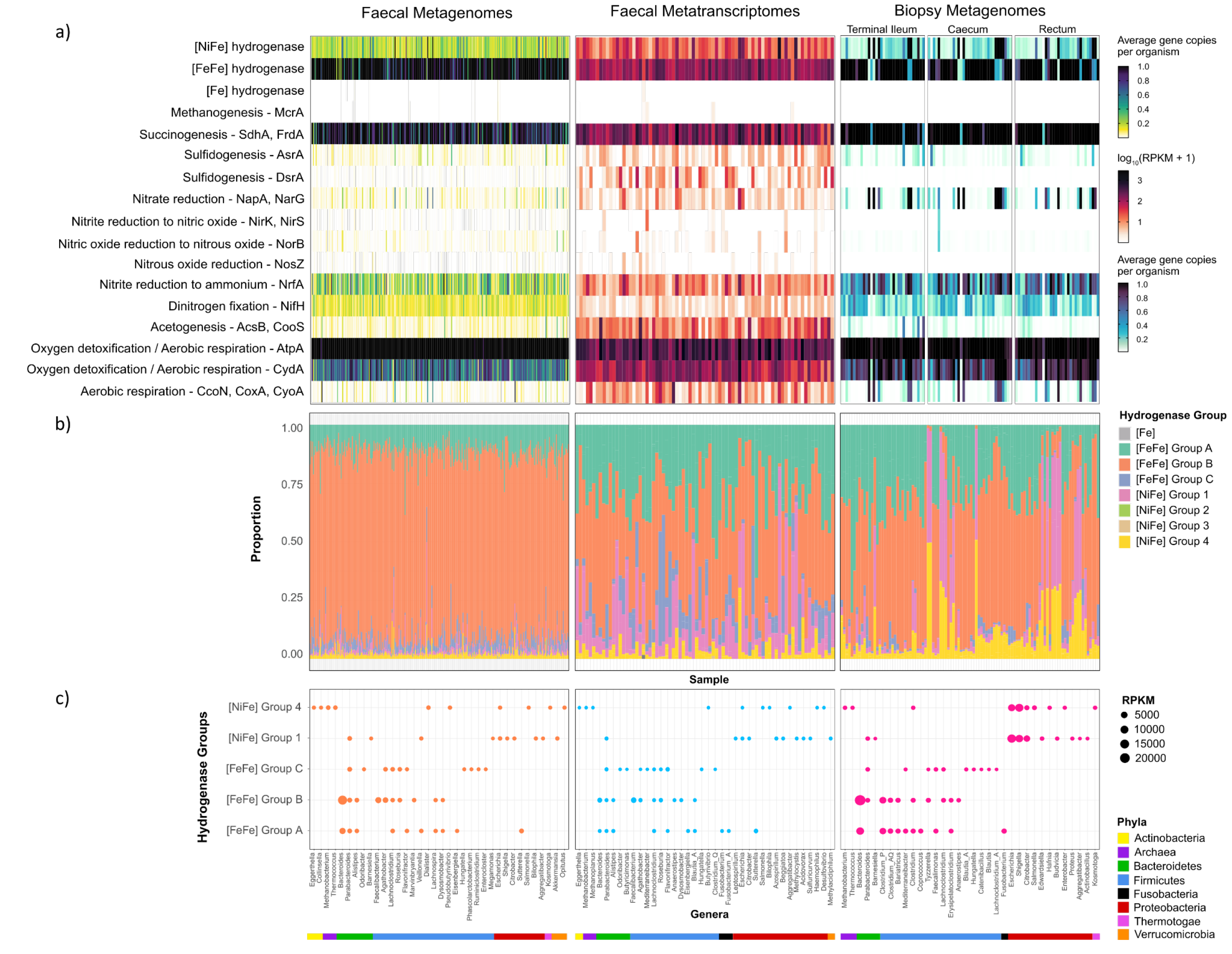
Abundance, expression, and distribution of hydrogenases and H₂-related metabolic genes throughout the human gut. (a) Abundance and expression of the genes encoding the catalytic subunits of the three types of hydrogenases and the terminal reductases known to use H_2_- derived electrons in faecal metagenomes (left; *n* = 300), faecal metatranscriptomes (middle; *n* = 78), and biopsy enrichment metagenomes (right; *n* = 102). These results summarise homology-based searches against comprehensive reference databases and are shown in average gene copies per organism (normalised to a set of universal single-copy ribosomal genes) for metagenomes and RPKM for metatranscriptomes. **(b)** Proportion of each hydrogenase group present in each sample per dataset. **(c)** Top genera predicted to encode or express hydrogenases for each dataset. The top 10 most abundant genera are included, for the five most abundant gut hydrogenase lineages, expressed in RPKM.

To infer which gut microbes encode these enzymes, we mapped the hydrogenase-encoding reads to both our comprehensive hydrogenase database (HydDB)^61^ and our in-house collection of 812 sequenced gut isolates **(Table S2)**. Group B [FeFe]-hydrogenases are very widespread among gut bacteria, encoded by 62% of isolates and the dominant gut phyla Firmicutes, Bacteroidetes, and Actinobacteria **(Table S2; Fig. 2)**. Their widespread conservation among Firmicutes and Bacteroidetes is demonstrated by the genome tree in **Fig. 2**. Based on read mapping, *Bacteroides* accounted for the most group B [FeFe]-hydrogenases in the metagenomes and metatranscriptomes, followed by *Alistipes* and Clostridia lineages such as *Faecalibacterium*, *Agathobacter*, and *Roseburia* **(Fig. 1c; Table S1)**. This finding suggests that this enzyme plays a core role in the lifestyles of diverse fermentative bacteria. The group A1 and A3 [FeFe]-hydrogenases were also widespread, encoded, and expressed by various Bacteroidia, Clostridia, and Fusobacteria genera, whereas formate hydrogenlyases were restricted to Enterobacteriaceae, Pasteurellaceae, and Coriobacteraceae **(Fig. 1c**; **Fig. 2; Table S1)**.

**Figure 2.**
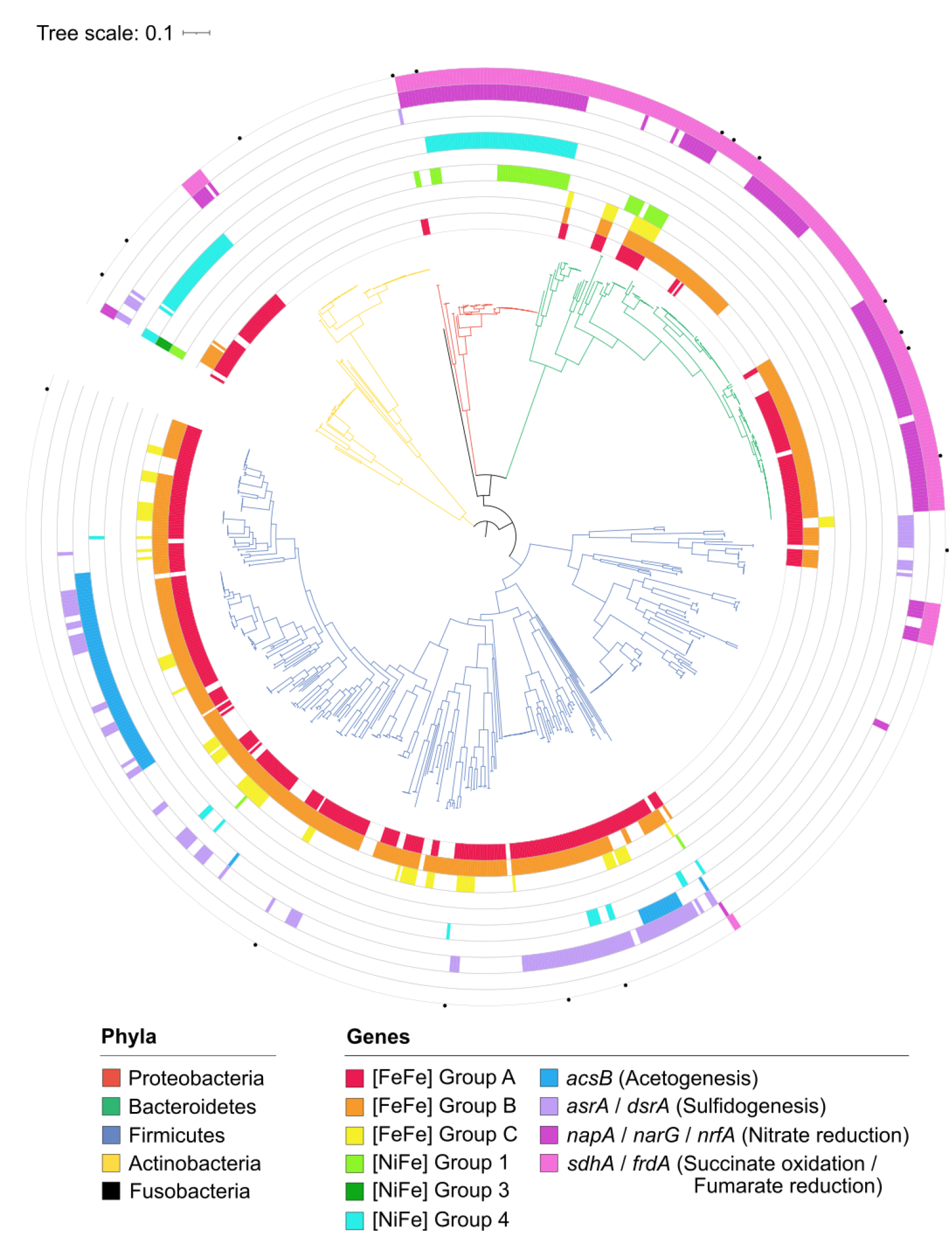
Phylogenomic tree showing distribution of hydrogenases among 812 bacterial isolates from the human gut. Isolates are from the five dominant phyla within the human gut with branch colours showing their phylum-level taxonomy. Isolates were shown to encode the catalytic subunit genes coding for the major groups of gut hydrogenases and the terminal reductases associated with methanogenesis, acetogenesis, sulfidogenesis, nitrate reduction, or succinogenesis (coloured rings). The tree was generated using approximately-maximum-likelihood estimation, Jukes-Cantor model (via FastTree) and the standardised ‘bac120’ phylogenetic analysis (*via* GTDB-Tk) and was midpoint rooted. Results are based on homology-based searches against comprehensive reference databases. Specific isolates were selected for further analysis, including culture-based activity measurements and transcriptome studies (black dots). Tree scale represents branch length of the tree, as calculated by number of base substitutions per base position.

A small but active proportion of the community are predicted to mediate H_2_ uptake in the human gut. Group 1 [NiFe]-hydrogenases, known to support anaerobic respiration using electron acceptors such as fumarate, nitrate, nitrite, and sulfite^37,38^, are encoded by 9% of gut bacteria based on metagenomic short reads **(Table S1; Fig. 1a & 1b)** and 6% of our isolates **(Table S2; Fig. 2)**. As evidenced by the extremely high standard deviations of their metagenome counts (0.09 ± 0.17 cpg) and metatranscriptome reads (51 ± 153 RPKM), these enzymes greatly vary between individuals **(Fig. 1a)**. They were primarily encoded and expressed by Enterobacteriaceae ([NiFe] group 1c, 1d), which are well known for using gut-derived H₂ as a respiratory energy source during colonisation^30,62,63^, as well as lineages such as *Veillonella* (1d), *Parabacteroides* (1d), and *Akkermansia* (1f) that remain to be investigated for their H₂ metabolism **(Fig. 1c)**. Some group A3 [FeFe]-hydrogenases were also encoded by hydrogenotrophic acetogens such as *Blautia*, where these enzymes oxidize H_2_, rather than produce it, in contrast to fermenters^64^. The group 3 and 4 [NiFe]-hydrogenases and [Fe]- hydrogenases of methanogenic archaea were also detected in a subset of samples. Consistently, we also detected genes encoding the signature enzymes responsible for fumarate, sulfite, nitrate, and nitrite reduction, acetogenesis, and methanogenesis in the metagenomes and metatranscriptomes **(Fig. 1a, Table S1)**. Importantly, although these enzymes except for fumarate reductase were in low abundance, they were often highly expressed (RNA / DNA ratio of 54 for acetyl-CoA synthase, 37 for dissimilatory sulfite reductase, 8 for periplasmic nitrate reductase) **(Table S1)**. Phylogenomic analysis of the gut isolates also revealed frequent co-occurrence of group 1 [NiFe]-hydrogenases with respiratory reductases **(Fig. 2)**. However, it should be noted that the respiratory reductases can accept electrons from a range of both organic and inorganic donors other than H_2_. Also detected were putative sensory hydrogenases (group C [FeFe]-hydrogenases, 0.11 ± 0.15 cpg) **(Fig. 1a)**, thought to differentially regulate [FeFe]-hydrogenases in response to H_2_ accumulation in Clostridia and likely other lineages^17,37,49^.

We tested whether these findings also extend to microbiota sampled within gut tissues, given stool samples provide a biased assessment of gut microbial content^65–67^. To do so, we collected mucosal biopsies from the terminal ileum, caecum, and rectum of 42 donors, then enriched and sequenced their microbiota^68^ **(Table S1)**. Concordantly group B [FeFe]-hydrogenases were by far the most abundant hydrogenases across these mucosal biopsy samples (0.75 ± 0.25 cpg); they were 3.7-fold more abundant than the next most abundant hydrogenase (group A3 [FeFe]-hydrogenase) and primarily encoded by *Bacteroides* based on read mapping **(Fig. 1a-c)**. The group 1c, 1d, and 4a [NiFe]-hydrogenases were also enriched by 6.1-, 2.6-, and 7.0-fold in the biopsy compared to stool metagenomes; this likely reflects the adherence of Enterobacteriaceae to the gut luminal walls, where they potentially use microbiota-derived H_2_ to support anaerobic and potentially even aerobic respiration **(Table S1; Fig. 1b & 1c)**. Thus, the group B [FeFe]-hydrogenase appears to drive fermentative H_2_ production throughout the intestines, much of which is likely recycled by respiratory hydrogenotrophs. No significant differences in hydrogenase content were found between intestinal regions, which was likely masked by the high degree of interindividual variation.

### Group B [FeFe]-hydrogenases are expressed and active in diverse gut isolates

While these analyses of metagenomes, metatranscriptomes, and isolate genomes respectively suggest the group B [FeFe]-hydrogenase is abundant, expressed, and widespread among gut bacteria, the precise activity of this enzyme remains unresolved. To confirm whether this enzyme is active, we used gas chromatography to test H_2_ production of 19 phylogenetically and physiologically diverse bacterial gut isolates each grown on standard YCFA medium under fermentative conditions **(Table S4; Fig. S1)**. Of these isolates, thirteen encoded group B [FeFe]-hydrogenases, either individually or together with other hydrogenases, all but one of which produced high levels of H_2_ **(Fig. 3a, Fig. S2)**. This collection included seven *Bacteroides* isolates that each rapidly produced headspace H_2_ to average maximum levels of 3.0 ± 0.6% during fermentative growth, as well as four genera from the class Clostridia **(Fig. 3b; Fig. S1; Table S3)**. We compared these activities to those of six control isolates that encoded either well-characterized lineages of H_2_-producing hydrogenases (group A1 [FeFe]- and group 4a [NiFe]-hydrogenases; positive controls) or lacked hydrogenases altogether (negative controls). The controls behaved as expected **(Fig. 3a; Fig. S1)**: no H_2_ was detected in the three isolates lacking hydrogenases (*Catenibacterium mitsuokai*, *Bifidobacterium longum*, and *Bacteroides stercoris*); high levels of H_2_ were produced during fermentative growth of a *Fusobacterium varium* isolate encoding prototypical group A1 [FeFe]-hydrogenase; and H_2_ was produced during fermentative survival in bacteria encoding the group 4a [NiFe]-hydrogenase containing formate hydrogenlyases (*Collinsella aerofaciens*, *Necropsobacter rosorum*) in line with their confirmed roles^42,69^. Altogether, these analyses show H_2_ production is a widespread trait among gut bacteria that encode group B [FeFe]-hydrogenases.

**Figure 3.**
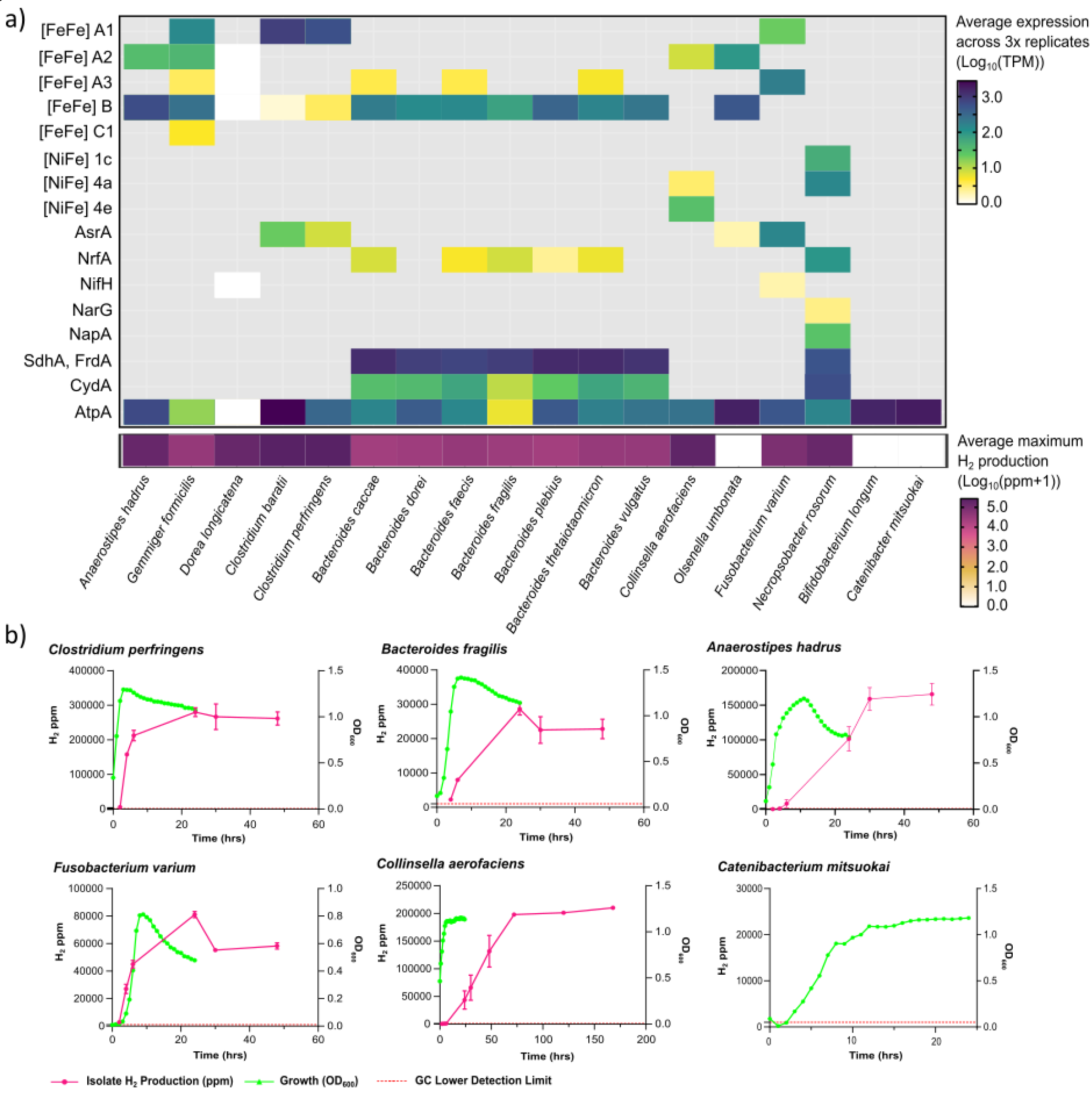
Hydrogenase expression and activity across 18 bacterial gut isolates. (a) The heatmap showing the average expression levels in transcripts per million (TPM) of the catalytic subunit genes for hydrogenases and the terminal reductases associated with sulfidogenesis, succinogenesis, nitrate reduction, and aerobic respiration. The bottom row shows the average maximum H_2_ production for each isolate. In both heatmaps, results show means from biologically independent triplicates. *B. longum* and *C. mitsuokai* do not encode hydrogenases and so are used as negative controls. **(b)** Bacterial growth measured by optical density (OD_600_, green lines) and H_2_ production (ppm; red lines), of representative isolates from chosen phyla over 24–168-hour periods (n = 3, mean ± SEM), where the lower detection threshold of the gas chromatograph is 1000 ppm (dashed red line).

We performed transcriptome sequencing to confirm whether the group B [FeFe]-hydrogenases are expressed and likely responsible for the observed activities **(Table S4)**. Patterns of hydrogenase expression and activity varied between species within the class Clostridia *Anaerostipes hadrus* encoded three [FeFe]-hydrogenases yet expressed the group B at higher levels (average 318 TPM) than its group A2 enzyme (33 TPM) **(Fig. 3a; Table S4).** Similarly, *Gemmiger formicilis* also exhibited higher expression of the group B hydrogenase (254 TPM) compared to its group A1 (139 TPM) and A2 (39 TPM) hydrogenases **(Fig. 3a; Table S4)**. These findings indicate that the group B [FeFe]- hydrogenase serves as the primary fermentative hydrogenase in both species. In contrast, the opportunistic pathogens *Clostridium perfringens* and *Clostridium baratii* expressed their prototypical fermentative group A1 [FeFe]-hydrogenases at much higher levels (*C. perfringens*: 305 TPM, *C.baratii*: 850 TPM) than their group B [FeFe]-hydrogenases (*C. perfringens:* 3.03 TPM, *C. baratii*: 0.42 TPM) **(Fig. 3a; Table S4)**. These extremely fast-growing bacteria also both produced much higher levels of H_2_ than the other isolates (up to 26.7% H_2_) **(Fig. 3a; Fig. S1)**. These findings are consistent with biochemical and genetic studies suggesting the group A1 enzyme predominates H_2_ production in *C. perfringens*^35,70^. Such *Clostridium* species appear to have evolved exceptionally rapid hydrogenases to enable vigorous growth in high nutrient conditions, though the metagenomic and metatranscriptomic analyses suggests they are in low abundance in most stool and biopsy samples **(Fig. 1)**. It remains unclear under which conditions such species express group B [FeFe]- hydrogenase. These findings suggest that species within the same phyla may employ distinct hydrogenases for similar purposes. *Dorea longicatena* generated substantial amounts of H₂, but transcriptomes yielded minimal reads mapping to metabolic genes and hence it is unclear whether its group B [FeFe]-hydrogenase is responsible **(Fig. 3a)**. In a further exception, the actinobacterium *Olsenella umbonata* did not produce detectable H_2_ despite encoding and expressing a group B [FeFe]-hydrogenase **(Fig. 3a)**. It is possible that its hydrogenase is active under specific conditions or alternatively this microbe internally recycles H_2_ using its group A2 [FeFe]-hydrogenase.

Our culture-dependent studies also indicated that fermentative H_2_ production is a key feature of *Bacteroides* physiology. After six hours of fermentative growth, the seven hydrogenase-encoding *Bacteroides* species had each produced between 0.68% and 2.50% levels of H_2_ in the headspace, averaging 1.51 ± 0.6% **(Fig. S1)**. For all seven strains, the group B [FeFe]-hydrogenase was expressed at high levels during growth, averaging 180 TPM (ranging from 71 ± 18 TPM for *B. fragilis* to 345 ± 17 TPM for *B. plebius*) **(Fig. 3a; Table S4)**. Four of these strains encoded the group B [FeFe]-hydrogenase as their sole H_2_-metabolizing enzyme. Three other strains (*B. caccae, B. thetaiotaomicron*, *B. faecis*) also encoded the trimeric electron-confurcating group A3 [FeFe]- hydrogenase, though the expression of this enzyme was minimal during these growth conditions (average 3.5 TPM) **(Fig. 3a; Table S4)**. Moreover, we observed no H_2_ production by *B. stercoris,* the only *Bacteroides* species in our isolate collection that consistently lacked any hydrogenase **(Fig. S2)**. These results show that the group B [FeFe]-hydrogenase accounts for the H_2_ production of *Bacteroides* during fermentative growth. In combination, the culture-dependent and culture- independent data suggest that these enzymes are highly conserved, expressed, and active across diverse gastrointestinal *Bacteroides* species. This suggests that this genus, despite not traditionally being associated with H_2_ cycling, is a dominant H_2_ producer in the human gut.

### *Bacteroides* use group B [FeFe]-hydrogenases to reoxidize ferredoxin during fermentation

We combined structural modelling, biochemical measurements, and metabolic reconstructions to confirm the activity and role of the group B [FeFe]-hydrogenase within *Bacteroides* **(Fig. 4)**. AlphaFold2 modelling **(Fig. S3 & S4)** confirms group B [FeFe]-hydrogenase are structurally conserved between *Bacteroides* species **(Fig. S5)** and form monomers with two distinct globular domains **(Fig. 4b)**: a H-cluster domain (containing the typical catalytic H_2_-binding H-cluster of [FeFe]- hydrogenases and two [4Fe4S] clusters) and a smaller ferredoxin-like domain (containing two [4Fe4S] clusters) connected through a short flexible linker. Consistent with being *bona fide* hydrogenases, these enzymes encode the three highly conserved sequence motifs of [FeFe]- hydrogenases, which line the binding pocket of the H-cluster (L1: T_294_SCCPSY_300_, L2: G_342_PCVAKRKE_350_, L3: E_448_VMACEGGCISGP_460_) **(Fig. 4b; Fig. S5)**; notably, Cys456 bridges the [4Fe4S] and catalytic di-iron site of the H-cluster, and Met450 coordinates the dithiolate bridgehead group of the di-iron site. Structural comparison with the well-characterized group A1 [FeFe]- hydrogenase (*C. pasteurianum* hydrogenase, CpI; PDB: 6GM2^71^) revealed that, while the H-cluster domain is largely conserved, the group B [FeFe]-hydrogenase is otherwise structurally unique (**Fig. 4a**). They particularly differ in their electron-relaying iron-sulfur clusters: whereas the group A enzyme contains a [2Fe2S] ferredoxin-like domain and a His-ligated [4Fe4S] cluster^72^, the group B enzyme is instead predicted to have a single ferredoxin-like domain containing 2×[4Fe4S] clusters. This ferredoxin-like domain is unusual in that its iron-sulfur clusters are distant from the main body of the enzyme as they are separated from the nearest [4Fe4S] cluster in the H-cluster domain by an edge-to-edge distance of at least 22 Å **(Fig. 4a)**. The distance between these two clusters is likely too far for effective electron transfer^73^, even after we accounted for conformational change driven by the flexible loops using AF-Cluster^74^. We hypothesise that the group B [FeFe]-hydrogenase interacts with soluble ferredoxins to enable productive electron transfer. Structural predictions also indicated that the group A3 [FeFe]-hydrogenases of *Bacteroides* are trimeric enzymes that confurcate electrons from reduced ferredoxin and NADH to H_2_ **(Fig. S6)**.

**Figure 4.**
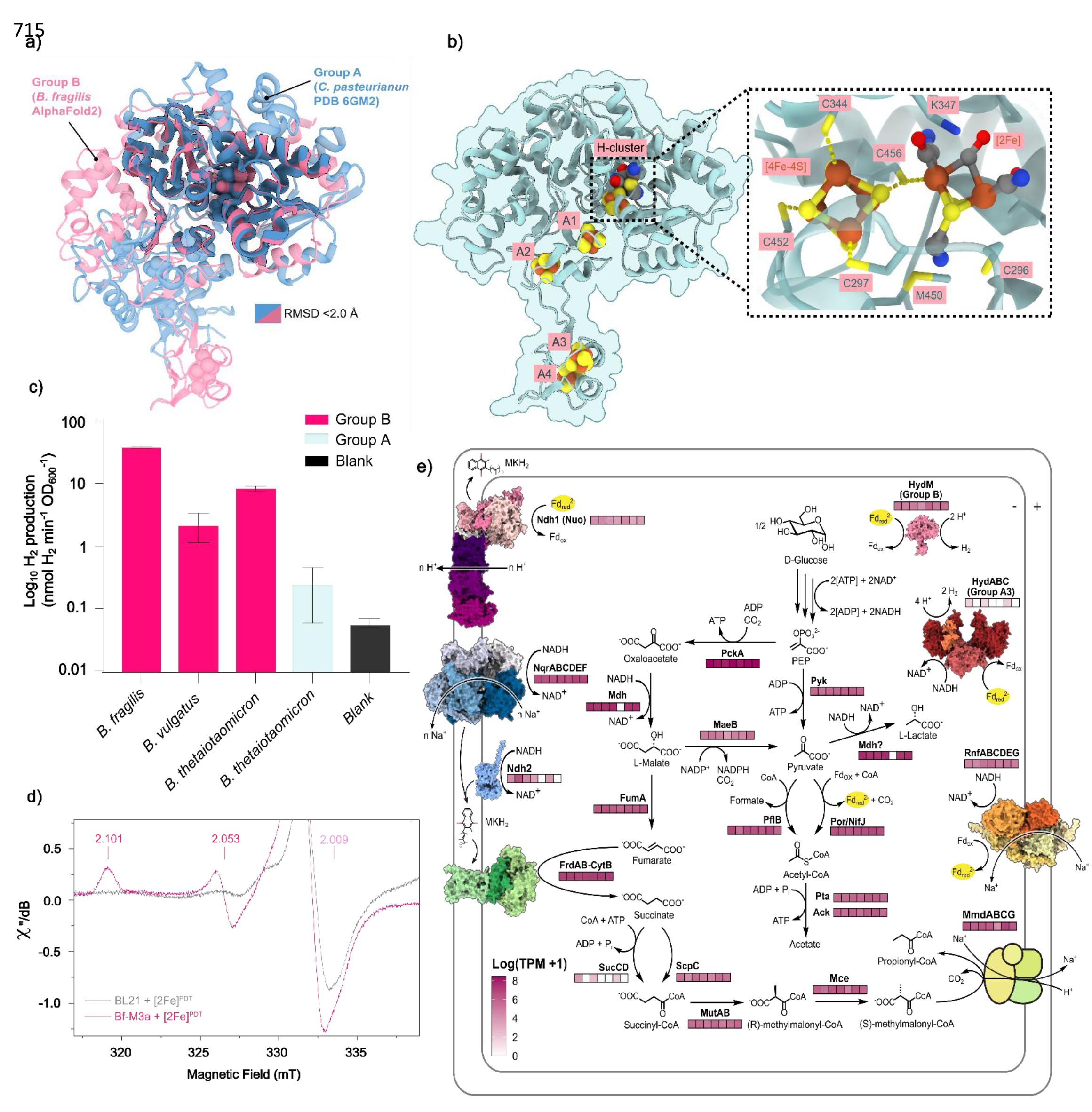
Metabolic integration, predicted structure, and biochemical activity of the group B [FeFe]-hydrogenases from *Bacteroides*. (a) A superposition of representative structures of the group A [FeFe]-hydrogenase (*Clostridium pasteurianum* CpI; X-ray crystallography; PDB: 6GM2^71^) and the group B [FeFe]-hydrogenase (*Bacteroides fragilis*; AlphaFold2). Portions where the two [FeFe]-hydrogenase groups show structural similarity (where RMSD <2 Å) are highlighted in bold and black outline. Divergence between the two structures (RMSD >2 Å) is depicted as transparent with no outline. RMSD: root mean square deviation. **(b)** Top-ranked predicted protein structure (AlphaFold2) of the *B. fragilis* group B [FeFe]-hydrogenase with putative [FeFe]-hydrogenase cofactors modelled. The predicted H- cluster site is shown in focus with conserved residues coordinating with the H-cluster labelled in green. Four iron-sulfur clusters are predicted to coordinate with conserved cysteines throughout the protein, labelled A1 to A4. **(c)** H_2_ production (measured by GC) monitored from cell lysates activated by addition of [2Fe]^adt^. Results are shown for the group B [FeFe]- hydrogenases of *B. fragilis*, *B. vulgatus*, and *B. thetaiotaomicron*, as well as the group A3 [FeFe]-hydrogenase of *B. thetaiotaomicron*. Activities were normalised for number of cells used (nmol H_2_ min^-1^ OD_600_^-1^) and error bars reflect standard deviation from biological triplicates. All enzymes were expressed in *E. coli* BL21(DE3) cells. The strain expressing the prototypical group A1 [FeFe]-hydrogenase from *Chlamydomonas reinhardtii* (*Cr*HydA1) was used as a positive control, while “Blank” represents the same strain but containing an empty vector that was also added with [2Fe]^adt^. **(d)** X-band EPR spectra recorded of cells expressing the *B. fragilis* group B [FeFe]-hydrogenase and empty vector BL21(DE3) control cells following anaerobic incubation with [2Fe]^pdt^. A distinct partial rhombic EPR signal attributable to the H_ox_ state of the H-cluster with the first two g-values (*g*_1_ = 2.101, *g*_2_ = 2.053) observable in [2Fe]^pdt^-treated hydrogenase-expressing cells, while the third *g*-value is not discernible due to signal overlap with cell background (see also Supplementary Note 1). EPR spectra were recorded at 20 K, 64 μW microwave power, and at a microwave frequency of 9.36 GHz. **(e)** Summary of the expression levels of fermentation genes in seven enteric *Bacteroides* isolates, including the group B and group A3 [FeFe]-hydrogenases. Expression is shown as TPM in boxes in the order of *B. caccae, B. faecis, B. fragilis, B. thetaiotaomicron, B. dorei, B. plebius, and B. vulgatus* under each relevant gene. Grey boxes indicate the gene is neither encoded nor expressed.

To determine whether *Bacteroides* [FeFe]-hydrogenases can bind the catalytic H-cluster and produce H_2_, we expressed their group B and A3 [FeFe]-hydrogenase catalytic subunit genes in *E. coli* BL21(DE3) cells **(Table S5; Fig. S7)**, activated lysates with the H-cluster mimic [2Fe]^adt^ as previously described^45,75,76^, and tested their ability to produce H_2_ using the standard methyl viologen as redox mediator and sodium dithionite as sacrificial electron donor. The group B [FeFe]- hydrogenases expressed from three different species all produced H_2_, in contrast to blank and empty vector controls, confirming that this group of enzymes are catalytically active **(Fig. 4c)**. Their relative activity varied compared to the positive control (the fast-acting group A1 [FeFe]-hydrogenase of *Chlamydomonas reinhardtii, Cr*HydA1^76–80^; at 30% (*B. fragilis*), 8% (*B. thetaiotaomicron*), and 2% (*B. vulgatus*) **(Fig. 4c; Table S5)**; these contrasting activities likely reflect differences in the expression, maturation, or solubility of these enzymes in the heterologous host, though are unlikely to be physiologically relevant given the species these enzymes were derived from each produced comparable amounts of H_2_ **(Fig. 3)**. In this recombinant system, the catalytic subunit of the group A3 [FeFe]-hydrogenase from *B. thetaiotaomicron* exhibited low activity close to the blank and empty vector controls when mixed with [2Fe]^adt^ **(Fig. 4c; Table S5)**.

Despite extensive effort, we were unable to purify stable or active group B [FeFe]-hydrogenases due to the low solubility of these enzymes. This prevented detailed comparisons of their kinetics, electrochemistry, or experimental structures compared to group A1 [FeFe]-hydrogenases. Nevertheless, we were able to demonstrate through whole-cell X-band EPR spectroscopy that the *B. fragilis* group B [FeFe]-hydrogenase, when incubated with the [2Fe]^pdt^ (a catalytically inactive cofactor mimic known to stabilise the di-iron site in a mixed valent Fe^I^Fe^II^ oxidation state such as EPR-active, H_ox_), produced spectroscopic signatures consistent with a typical H-cluster^75,81^ **(Fig. 4d)**. Cell suspensions displayed a partially resolved rhombic EPR signal with *g*_1_ = 2.101 and *g*_2_ 2.053 (**Fig. 4d**). The observed *g*-values suggest formation of the H-cluster in an H_ox_-like state and support the notion that the H-cluster of the group B [FeFe]-hydrogenase has an electronic structure similar to the distantly related prototypical group A enzymes **(Table S6)**. As elaborated in **Supplementary Note 1**, the third *g*-value (*g*_3_) was not observed likely due to being obscured by unrelated signals. In combination, the structural predictions and recombinant analysis suggest the group B [FeFe]- hydrogenases are true hydrogenases that bind the H-cluster and produce H_2_, though differ from other hydrogenases in their redox centres and electron flow pathways.

We used the transcriptomes of the seven *Bacteroides* species to predict their central carbon metabolism and infer how their [FeFe]-hydrogenases likely integrate **(Table S3)**. Supporting previous physiological observations, these reconstructions suggest that all species are mixed-acid fermenters^82–85^ that can break down sugars to pyruvate through the glycolysis pathway, convert pyruvate to acetate (*via* pyruvate-ferredoxin oxidoreductase and acetate kinase); and also reduce oxaloacetate to succinate (*via* enzymes including fumarate reductase) and propionate (*via* methylmalonyl-CoA pathway). Consistently, these bacteria all express the genes for these pathways at similarly high levels during mid-exponential fermentative growth **(Fig. 4e)**. The group B [FeFe]- hydrogenase likely primarily reoxidizes the ferredoxin reduced by the pyruvate-ferredoxin oxidoreductase (PFOR) during acetate production, disposing these excess electrons as H_2_. The Rnf complex, which couples Na^+^/H^+^-import to reverse electron transport from NADH to oxidized ferredoxin, is potentially an additional source of reduced ferredoxin for the hydrogenase^86,87^; however, this physiologically reversible complex may also serve as a reduced ferredoxin sink that generates sodium/proton-motive force **(Fig. 4e)**. All other reductive branches of the *Bacteroides* fermentation pathways, namely for succinate, propionate, and lactate formation, do not directly compete with H_2_ production via the Group B [FeFe]-hydrogenase for electrons, but will indirectly reduce H_2_ production by shunting pyruvate away from PFOR. Pyruvate-formate lyase provides an additional PFOR bypass via the redox-neutral production of formate and acetyl-CoA **(Fig. 4e)**. Whereas the genes for the succinate/propionate branch and PFOR were relatively consistently expressed across the strains, expression of pyruvate-formate lyase and a putative lactate dehydrogenase varied as much as 4.5- and 50-fold respectively **(Table S3)**; thus, lactate and formate production may be the most important alternate routes of electron disposal that compete with H_2_ production in *Bacteroides* species. The group A3 [FeFe]-hydrogenase potentially contributes to redox homeostasis, likely by coupling oxidation of reduced ferredoxin and NADH to H_2_ production, and may be important under certain conditions **(Fig. 4e)**; however, during fermentative growth, the enzyme remains expressed at low levels, likely reflecting that NADH consumption by this enzyme would compete with the reductant used for fumarate and propionate production and thereby ATP production through substrate-level phosphorylation.

### Group B [FeFe]-hydrogenases are depleted in gastrointestinal disorders and other diseases

To investigate the links between hydrogen metabolism with health and disease, we compared the levels of hydrogenase-associated genes based on stool metagenomes of 871 healthy individuals and 790 diseased individuals, based on a case-control study of Crohn’s disease (CD)^88^ and reports on 11 other chronic disease phenotypes^55^ **(Fig. 5)**. Consistent with the above analyses **(Fig. 1)**, group B [FeFe]-hydrogenases were the most abundant hydrogenase genes overall, though their levels often varied between individuals **(Fig. 5; Table S7)**. However, their average levels were significantly higher (*p =* 0.0023) in healthy individuals (0.72 ± 0.17 cpg) compared to those with Crohn’s disease (0.56 ± 0.24 cpg). Contrastingly, there were strong increases in the average levels of the prototypical fermentative hydrogenase (group A1, 2.8-fold, *p* = 6.6 × 10^-7^), formate hydrogenlyase (group 4a, 5.2-fold, *p* = 6.8 × 10^-6^), and to lesser extent the electron-confurcating hydrogenase (group A3, 1.4-fold, *p* = 0.04) in Crohn’s disease individuals. Most remarkably, the ratio of group B to group A1 [FeFe]-hydrogenases shifted by 2.3-fold between the two cohorts (*p* = 1.9 × 10^-11^). Capacity for H_2_ oxidation also increased, with a 2.6-fold increase in respiratory group 1d [NiFe]-hydrogenase genes in Crohn’s disease (*p* = 3.8 × 10^-5^) **(Fig. 5)**. Though these differences may be potentially only correlative, altered H_2_ cycling may contribute to the Crohn’s disease phenotype through various possible mechanisms. For example, intestinal respiratory bacteria (e.g. Enterobacteriaceae) may benefit from elevated H_2_ production by the highly active group A1 [FeFe]- hydrogenases, by using this electron donor to reduce inflammation-derived electron acceptors. Consistently, a previous study showed increased H_2_ oxidation contributes to the expansion of *E. coli* during gut inflammation in a murine model^31^. There was also a significant enrichment of group A1 compared to group B [FeFe]-hydrogenases in several other chronic disease states, including atherosclerosis, liver cirrhosis, colorectal cancer, and type 2 diabetes **(Fig. S8)**. A range of other significant variations were also observed, including a near-absence of group 1d [NiFe]- hydrogenases in type 2 diabetes (*p* = 7.8 × 10^-9^) **(Fig. S9)**. Further mechanistic studies are required to better understand the basis of these differences.

**Figure 5.**
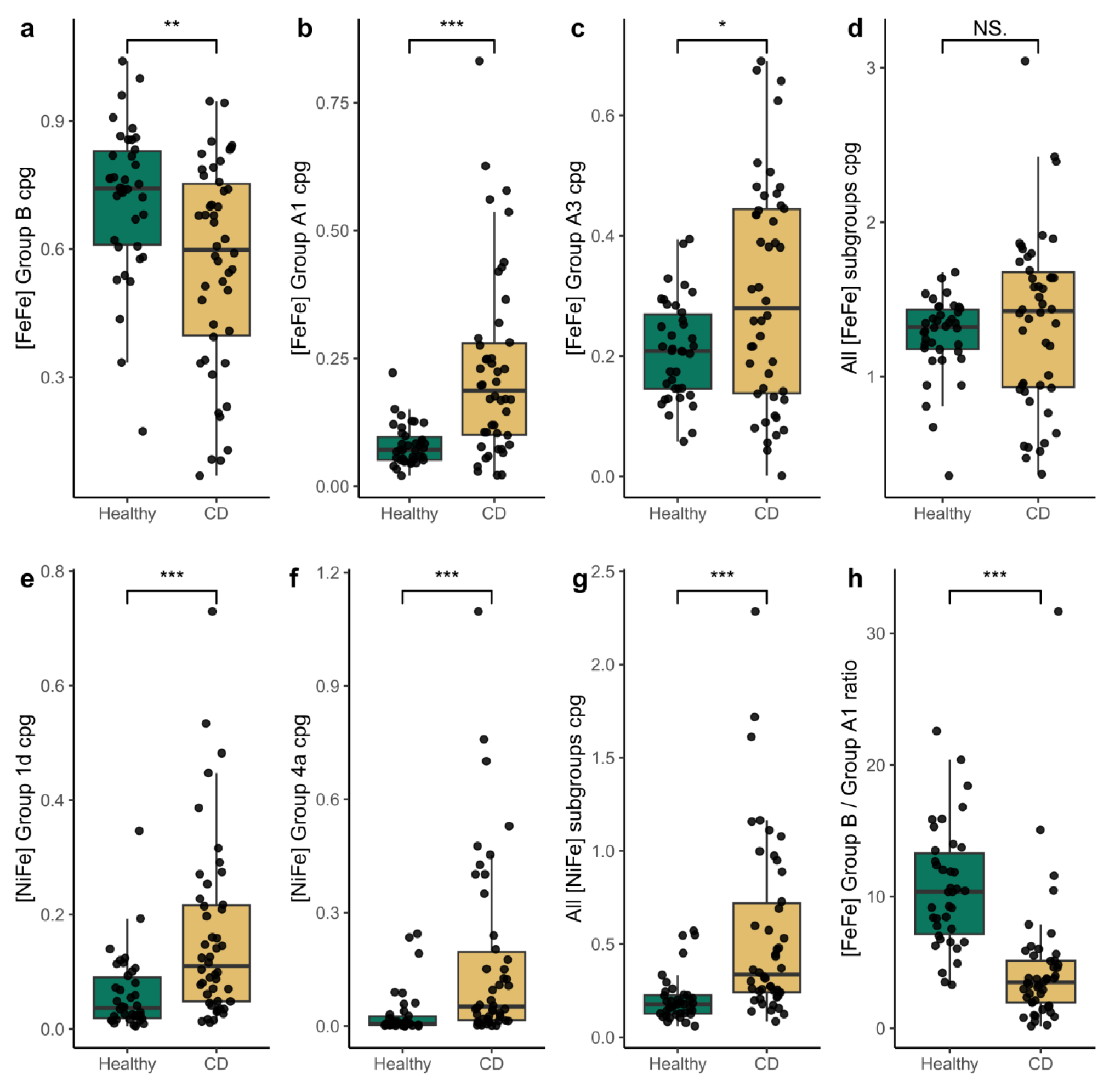
Comparison of hydrogenase gene levels in healthy individuals compared to Crohn’s disease (CD) patients in a case-control study. Sum of counts per genome are shown for **(a)** group B [FeFe]-hydrogenases, **(b)** group A1 [FeFe]-hydrogenases, **(c)** group A3 [FeFe]-hydrogenases, **(d)** all [FeFe]-hydrogenase subgroups, **(e)** group 1d [NiFe]-hydrogenases, **(f)** group 4a [NiFe]- hydrogenases, and **(g)** all [NiFe]-hydrogenase subgroups. Also shown is **(h)** the ratio of group B to group A1 [FeFe]-hydrogenases. Statistical significance was assessed with Wilcoxon tests, where * *p* < 0.05, ** *p* < 0.01, *** *p* < 0.001. Box plots show median (centre line), upper and lower quartiles (box limits), 1.5 × interquartile range (whiskers), and individual samples. n = 46 Crohn’s disease patients, n = 38 healthy controls.

## Conclusions

By integrating analyses at the enzyme, cellular, and gut ecosystem levels, we provide multifaceted evidence that the group B [FeFe]-hydrogenase mediates H_2_ production in diverse bacteria and drives fermentation in the healthy human gut. These observations also suggest that *Bacteroides*, a genus previously unrecognised as major H₂ producers, plays a more central role in gut H₂ cycling than initially understood and uses a hydrogenase of previously unknown function. It remains unclear what competitive advantage is conferred by the group B [FeFe]-hydrogenase compared to the functionally similar group A1 [FeFe]-hydrogenase. Both enzymes are predicted to be monomeric ferredoxin- dependent H_2_-producing enzymes with similar active site structures and biosynthetic pathways. Nevertheless, the group B enzyme is unique for its ferredoxin-like domain separated by a flexible linker and also seems to have somewhat lower activity than its group A1 counterparts based on the cellular and whole-cell data. Detailed side-by-side studies of the protein-protein interactions, kinetics, electrochemistry, and oxygen sensitivity of the purified enzymes may help disentangle their differences. Nevertheless, it is apparent that the group B enzyme has been selected in diverse bacterial species to produce high levels of H_2_ during fermentative growth^44^. This hydrogenase is particularly abundant in healthy people and may be an indicator of H_2_ homeostasis, whereas there is a shift in favour of group A1 [FeFe]-hydrogenases in disease states such as Crohn’s disease.

This study also provides a holistic perspective on the microbes and enzymes responsible for H_2_ cycling in the human gut. We provide the most in-depth study of the distribution of gastrointestinal hydrogenases to date, surpassing our last bioinformatics survey in this area that was limited to just 20 metagenomes, and bridge genomic insights with culture- and enzyme-based validation^19^. We show H_2_ production is an extremely widespread trait, demonstrating this trait extends to gut Bacteroidetes, Fusobacteria, Actinobacteria and Proteobacteria in addition to the well-studied Clostridia. Our findings also highlight that the mediators of the three conventionally described pathways for H_2_ disposal, namely methanogenesis, acetogenesis, and sulfidogenesis, are in low abundance but are transcriptionally active in the gut^3,12,14^. Given their high abundance in both stool and biopsy metagenomes, it is nevertheless likely that respiratory bacteria that use electron acceptors such as fumarate, nitrate, and sulfoxides are also major and potentially dominant hydrogenotrophs, especially the Enterobacteriaceae. These lineages are especially enriched in certain disease states, such as Crohn’s disease, where they may support both aerobic and anaerobic respiration using inflammation-derived electron acceptors. Follow-up studies should combine metagenomic, biochemical, and culture-based studies to determine which processes and microbes dominate H_2_ oxidation in the human gut. Further studies are also required to better characterise the roles of some of the moderately abundant but still functionally characterised hydrogenases identified here, including the group A2 [FeFe]-hydrogenases, group 4e [NiFe]-hydrogenases, and the sensory hydrogenases. There is also critical need to better understand what drives the vast interindividual variation in hydrogenase composition and expression between individuals, and how this relates with gastrointestinal function and disease states. In summary, our multifaceted approach uncovers the abundance, diversity, and functional roles of hydrogenases of previously unrecognised importance in the human gut during health and disease.

## Materials and methods

### Mucosal biopsy sampling and metagenomes

Mucosal biopsy metagenome samples were obtained from 102 mucosal biopsy enrichment metagenomes of 42 paediatric patients receiving colonoscopy due to non-inflammatory conditions at the Monash Children’s Hospital (Monash Health; Victoria, Australia; Human Research Ethics Committee (HREC) (HREC/16/MonH/253) and Monash University Ethics Committee (Monash Health ref. 16367A). Samples were obtained from the terminal ileum, caecum and rectum and transferred to anaerobic conditions within 15 minutes of collections. Biopsy metascrapes were performed after 24 hour incubation on YCFA agar plates at 37°C under anaerobic as described previously^68^ with resulting DNA extracted using the MP Biomedicals FastDNA SPIN Kit for soil and sequenced on the Illumina NextSeq2000. Resulting data is accessible via the European Nucleotide Archive (ENA) under accession number PRJEB45397 (https://www.ebi.ac.uk/ena/browser/view/PRJEB45397).

### Stool metagenome and metatranscriptome datasets

For the stool metagenome and metatranscriptome analyses, raw paired-end short reads were obtained from an IBD microbiome functionality study^57^, accessed via the European Nucleotide Archive (ENA) under accession number PRJNA389280 (https://www.ebi.ac.uk/ena/browser/view/PRJNA389280). The dataset included 78 paired metagenomes and metatranscriptomes, along with 222 additional metagenomes from the faecal samples of 117 healthy patients as well as those with varying gastrointestinal IBD pathologies.

### Metagenomic and metatranscriptomic analyses

For the stool metagenome and metatranscriptome analyses, raw paired-end short reads were obtained from an IBD microbiome functionality study^57^, accessed via the European Nucleotide Archive (ENA) under accession number PRJNA389280 (https://www.ebi.ac.uk/ena/browser/view/PRJNA389280). The dataset included 78 paired metagenomes and metatranscriptomes, along with 222 additional metagenomes from the faecal samples of 117 patients. The stool metagenome, stool metatranscriptome, and mucosal biopsy metagenomes were quality-checked using FastQC (v0.11.7)^89^ and MultiQC (v1.0)^90^. Adapter and PhiX sequences, and low-quality bases were trimmed and filtered with BBDuk from BBTools (v38.51) suite^91^. SortmeRNA (v4.3.3)^92^ removed rRNA sequences from metatranscriptomic reads. Resulting cleaned forward reads were screened (blastx) using DIAMOND v2.0.^93^ against the HydDB dataset^61^ and a manually curated in-house database including enzymes associated with H_2_- producing and H_2_-consuming pathways (https://doi.org/10.26180/c.5230745). Alignments were filtered to a minimum length of 28 amino acids, and further based on minimum percentage identity thresholds previously determined and validated for each protein in the database: 50% (AcsB, ArsC, AsrA, CcoN, CooS, CoxA, CydA, CyoA, DsrA, FdhA, NapA, NarG, NiFe (60% for group 4), NifH, NirK, NorB, NosZ, NrfA, RHO, SdhA_FrdA and Sqr), 60% (FeFe and NuoF) and 70% (AtpA and YgfK). Read counts were normalised to reads per kilobase million (RPKM), and metagenomes were further normalised against mean RPKM values estimated from 14 single-copy ribosomal marker genes to obtain an ‘average gene copy per organism’ value for each gene. For predicted hydrogenase sequence reads, the taxonomy of the best hit was retrieved and summarised in RPKM to evaluate which taxonomic groups contribute most of these reads.

### Gut isolate genomic analysis

Whole genome sequences of 818 gut isolates from adult and paediatric faecal and biopsy samples were obtained from the Australian Microbiome Culture Collection (AusMicc; https://ausmicc.org.au/) and a previous study describing a collection of gut isolate genomes (HBC)^94^. Resulting genomes were quality checked with CheckM (v1.1.3)^95^ and those with >90% completeness and <5% contamination (n = 812) were retained. Protein sequences of the retained genomes were used to search for and identify alignments matching the previously mentioned protein database using the blastp function of DIAMOND (v2.0.9)^93^. Alignment criteria included query and subject coverage thresholds set at 80%, and further filtering was conducted based on the previously mentioned percentage identity thresholds for each database protein. GTDB- Tk (v1.6.0) (database R06-RS202)^96^ was used to assign a taxonomic classification to each isolate using the “classify_wf” option, and a phylogenetic tree was constructed with the “de_novo_wf” option. The tree was visualized, midpoint-rooted, and the copy number per isolate genome of relevant hydrogen-related metabolic genes and hydrogenase subgroups was overlaid using the Interactive Tree of Life (iTOL)^97^ to observe differences in hydrogen metabolism across the different phylogenetic groups.

### Bacterial growth analyses

All isolates used in this study, sourced from healthy human faecal samples, were obtained from AusMiCC^68,94^. Nineteen isolates were selected to compare the expression and activity of the group B [FeFe]-hydrogenases compared to other H_2_-producing bacteria across taxonomically diverse gut bacteria, including three control isolates lacking hydrogenases and three positive controls encoding well-characterised H_2_ producing hydrogenases^42,69^. **Table S3** lists the strains and their hydrogenase content. All isolates were accessed from glycerol stocks containing Yeast Casitone Fatty Acids (YCFA) broth media^98^ with 25% glycerol, stored at -80°C, and revived in pre-reduced YCFA broth. Incubation was carried out anaerobically at 37°C in an atmosphere of 10% H₂, 10% CO₂, 80% N₂ for 24 hrs. Solid media, when required, was supplemented with 0.8% w/v of bacterial agar. Growth assessment involved measuring the optical density (OD_600_) of each isolate over 24 hours while anaerobically cultured in YCFA broth at 37°C. Each isolate, in duplicate, was sub-cultured into a 200 µl 96-well plate, with a 1:100 dilution of culture to broth. An hourly assessment of OD_600_ was conducted using a FLUOstar Omega Microplate Reader, with readings taken under anaerobic conditions, and shaking before each measurement.

### Hydrogen production assays

For the H_2_ production assay, isolates were plated on pre-reduced YCFA agar plates and grown anaerobically at 37°C for 24 hrs. A single colony was used to inoculate 3 mL of pre-reduced YCFA broth in a 15 mL Falcon tube, which was incubated anaerobically at 37°C for 24 hrs. After incubation, each starter culture was used to inoculate triplicate 30 mL aliquots of YCFA broth to a starting OD_600_ of 0.025. Cultures were maintained in 120 mL glass serum vials sealed with lab-grade butyl rubber stoppers. Immediately after inoculation, the headspace of each culture vial was flushed for 10 minutes with 99.99% pure N₂ to remove residual H_2_ and ensure that production of H_2_ was thermodynamically favourable, and entirely biotic in origin. Gas chromatography was used to assess the H₂ production capabilities of each isolate over time. To establish a baseline H₂ concentration for each isolate (in triplicate), a gas-tight syringe was used to collect initial headspace gas samples from each culture immediately after N₂ flushing. Headspace gas samples were then collected at pre-determined time points based on growth curve data, and until increases in H₂ concentration were no longer detected. H₂ concentration was measured using a gas chromatograph containing a pulse discharge helium ionisation detector (model TGA-6791-W- 4U-2, Valco Instruments Company Inc) as previously described ^99^. This gas chromatograph was able to detect a wide range of H₂ concentrations (0.1% – 10% H₂), however, sample dilution of 2.5× was necessary to measure the H₂ produced by the isolates within the quantifiable range. Calibration samples of known H₂ concentration were used to quantify H_2_ in parts per million. The H₂ concentration within the media-only control vials was measured concurrently to confirm that H₂ production in isolate samples was biotic.

### RNA extraction

Transcriptomic analysis was performed for all isolates to verify the active expression of hydrogenases identified within the genome. Triplicate cultures of each isolate were grown under the same conditions as described for the H_2_ production assay. Cells were harvested for RNA extraction during active H₂ production at either exponential phase (isolates with a group A or B [FeFe]-hydrogenase) or stationary phase (isolates with a group 4a [NiFe]-hydrogenase), as indicated by previously conducted growth curves. To quench cells, a glycerol-saline solution (3:2 v/v, -20°C) was added prior to centrifugation (4500 × *g*, 30 min, -9°C). The cell pellet was resuspended in 1 mL of an additional glycerol-saline solution (1:1 v/v, -20°C) and centrifuged again (4,500 × *g*, 30 min, -9°C). Cell pellets were then resuspended in 1 mL TRIzol reagent, transferred to a tube containing 0.3 g of 0.1 mm zircon beads, and subjected to five cycles of bead-beating (30 seconds per cycle, 5000 rpm, resting on ice for 30 seconds between cycles) using a Bertin Technologies ‘Precellys 24’ bead-beater before centrifugation (12,000 × *g*, 10 minutes at 4°C). Supernatant was transferred to a new tube and 200 µl of chloroform was added, inverted to mix for 15 seconds, then incubated at room temperature for 2-3 minutes prior to centrifugation (10,000 × *g*, 15 minutes at 4°C) for phase separation. The aqueous phase underwent purification using the RNeasy Mini Kit following the manufacturer’s instructions (QIAGEN), with on-column DNA digestion using the RNase-free DNase Kit (RNeasy Mini Handbook, QIAGEN). RNA was eluted into RNase-free water, and the concentration for each sample was determined using the RNA HS Qubit Assay Kit according to manufacturer’s instruction (Thermo Fisher Scientific).

### Transcriptome sequencing

The Monash Health Translation Precinct Medical Genomics Facility prepared libraries using the Illumina ‘Stranded Total RNA prep with Ribo-Zero plus Microbiome’ kit. A total of 200 ng of RNA underwent 16 cycles of amplification. Final libraries were quantified by Qubit, combined into an equimolar pool, and quality-checked by Qubit, Bioanalyzer, and qPCR. For sequencing, 1000 pM of the library pool was clustered on a P2 NextSeq2000 run and 59 bp sequencing was performed. The total run yield was 66.56 G, with approximately 496.7 million reads passing filter, achieving a %Q30 of 92.57. Transcriptomic data was quality checked and pre- processed using FastQC (v0.11.7)^89^, MultiQC (v1.0)^90^ and BBDuk from BBTools suite (v38.51)^91^ as above. Successful ribodepletion was confirmed by SortMeRNA (v4.3.3)^92^. Each isolate’s genome was annotated using Prokka (v1.14.6)^100^, and transcript expression was quantified by mapping the transcripts to these annotated genomic features using Salmon (v1.9.0)^101^ with default settings (salmon quant). Gene expression was quantified as relative abundance in transcripts per million (TPM). To identify transcripts matching previously identified hydrogenase hits, Prokka-generated annotated protein sequence files were validated with DIAMOND alignment as described above. Transcript IDs were used to match the hydrogenase hits for each isolate to the corresponding TPM values, for evaluation of hydrogenase expression. For metabolic pathway analysis, DRAM (v.1.4.6)^102^ was used to annotate each transcriptome with the KEGG protein database^103^. The genome of *B. fragilis* was incomplete and lacking mapped *atpA* and *cydA* genes, so the genome of a reference strain from NCBI (ASM1688992v1) was used to map these genes to and demonstrate their expression.

### AlphaFold2 structural modelling

Protein structure predictions from *Bacteroides* Group B [FeFe]- hydrogenase sequences **(Table S4)** were generated using AlphaFold2 (v2.1.1)^104,105^ through the ColabFold (v1.5.2)^106^ notebook. The specified ColabFold parameters were as follows: num_relax (1), template_mode (none), msa_mode (mmseqs2_uniref_env), pair_mode (unpaired_paired), model_type (alphafold2_ptm), pairing_strategy (greedy). For the *B. fragilis* group B [FeFe]- hydrogenase model (*Bf*HydM), num_recycles was set to 48, whereas *B. thetaiotaomicron* and *B. vulgatus* num_recycles were set to 3. For the *B. thetaiotaomicron* group A3 [FeFe]-hydrogenase model (*Bt*HydABC), num_recycles was set to 48. To model cofactors into the predicted *Bf*HydM and *Bt*HydABC apo structures, the Foldseek^107^ web server was used to search the PDB100 database for experimental structures with similar folds to *Bf*HydM and *Bt*HydABC. The following Foldseek parameters were used: databases (PDB100 2201222), mode (3Di/AA), taxonomic filter (none). For *Bf*HydM, two experimental structures returned by Foldseek, PDB 8ALN^108^ and 1FCA^109^, exhibited high structural similarity to the input, while also containing iron-sulfur clusters and a H-cluster **(Fig. S4)**. Similarly, for *Bt*HydABC, three experimental structures returned by Foldseek were used for cofactor modelling, PDB 8A5E^64^, 1FEH^40^, and 1FCA^109^ **(Fig. S6)**. UCSF ChimeraX (v1.6.1)^110^ was used to align these experimental structures to the predicted *Bf*HydM and *Bt*HydABC models with the matchmaker command (Needleman-Wunsch algorithm setting). Cofactors were added in corresponding positions to those of the experimental structures as shown in **Fig. S3** and **Fig S6**. At sites where the AlphaFold2 model and the experimental structures differed, cofactors were manually positioned and adjusted to optimise coordination and to minimise clashes, and bond lengths were assessed to ensure they were biochemically valid.

### Protein expression and preparation

Chemicals used for protein production and characterisation were purchased from VWR and used as received unless otherwise stated. Genes encoding the group B [FeFe]-hydrogenases of *B. fragilis*, *B. vulgatus*, and *B. thetaiotaomicron* and group A3 [FeFe]-hydrogenase of *B. thetaiotaomicron* **(Table S5)** were cloned into pET-11a(+) by Genscript, using restriction sites *Nde*I and *Bam*HI following codon optimisation for expression in *Escherichia coli*. Chemically competent *E. coli* BL21(DE3) cells were transformed using the constructs to express the apo-forms of the hydrogenases lacking the diiron subsite of the H-cluster. Starter cultures were grown overnight in 5 mL LB medium containing 100 µg mL^-1^ ampicillin at 37°C. These cultures were subsequently used to inoculate 80 mL of M9 medium (22 mM Na_2_HPO_4_, 22 mM KH_2_PO_4_, 85 mM NaCl, 18 mM NH_4_Cl, 0.2 mM MgSO_4_, 0.1 mM CaCl_2_, 0.4% (v/v) glucose) containing 100 µg mL^-1^ ampicillin. Cultures were grown at 37°C and 150 rpm until reaching an optical density (OD_600_) of approximately 0.4 to 0.6. Protein expression was induced by the addition of 0.1 mM FeSO_4_ and 1 mM IPTG. Induced cultures were incubated at 20°C and 150 rpm for approximately 16 h. Cells were thereafter harvested by centrifugation at 4,930 × *g* for 10 mins at 4°C. All subsequent operations were carried out under anaerobic conditions to prevent hydrogenase inactivation by atmospheric oxygen in an MBRAUN glovebox ([O_2_] < 5 ppm). The cell pellet was resuspended in a 0.5 mL lysis buffer (30 mM Tris-HC pH 8.0, 0.2 % (v/v) Triton X-100, 0.6 mg mL^-1^ lysozyme, 0.1 mg mL^-1^ DNase, 0.1 mg mL^-1^ RNase). Cell lysis involved three cycles of freezing/thawing in liquid N_2,_ and the supernatant was recovered by centrifugation (29,080 × *g*, 10 mins, 4°C).

### H_2_ production assays of activated hydrogenases

The H_2_ production assays followed established protocols with minor modifications^76^. In short, the [2Fe]_H_ subsite mimic, (Et_4_N)_2_[Fe_2_(µ- SCH_2_NHCH_2_S)(CO)_4_(CN)_2_] ([2Fe]^adt^), was synthesised in accordance to previous protocols with minor modifications and verified by Fourier transform infrared (FTIR) spectroscopy (Li and Rauchfuss, 2002; Zaffaroni *et al.*, 2012). Incorporation of cofactor involved the addition of 100 µg of the [2Fe]^adt^ subsite mimic (final concentration 80 μM) to 380 μL of the supernatant in potassium phosphate buffer (100 mM, pH 6.8) and 1 % (v/v) Triton X-100. The reaction mixture was anaerobically incubated at 20°C for 1-4 hr in a sealed vial. The non-purified lysate containing the [2Fe]^adt^ subsite mimic was mixed with 200 μL of potassium phosphate buffer (100 mM, pH 6.8) with 10 mM methyl viologen and 20 mM sodium dithionite. Reactions were incubated at 37°C for up to 120 mins. H_2_ production was determined by analysing the reaction headspace after 15 mins using a PerkinElmer Clarus 500 gas chromatograph (GC) equipped with a thermal conductivity detector (TCD) and a stainless-steel column packed with Molecular Sieve (60/80 mesh). The operational temperatures of the injection port, oven, and detector were 100°C, 80°C, and 100°C, respectively. Argon was used as carrier gas at a flow rate of 35 mL min^−1^. The strain expressing prototypical *Cr*HydA1^76–80^ served as a positive control, while “Blank” denoted the same strain, but containing an empty vector that was also added with [2Fe]^adt^. Three biological replicates were run at varying times (1-4 hours) of incubating the cell lysates with the [2Fe]^adt^ subsite mimic. Incubation time was not found to influence the observed H_2_ production. Thus, variation in H-cluster formation rates did not appear to have a substantial influence on the outcome of the screening process.

### Whole-cell EPR spectroscopy

Samples for whole-cell electron paramagnetic resonance (EPR) spectroscopy were prepared following a previously published protocol with minor modifications^76^.The cell pellet from 80 mL cultures (see Protein expression and preparation) was resuspended in 1 mL M9 medium, flushed with N_2_ gas for 10 mins, and mixed with a [2Fe]_H_ subsite mimic that lacks the natural nitrogen bridgehead of [2Fe]^adt^ to propane-1,3-dithiolate ([2Fe]^pdt^, (Et_4_N)_2_[Fe_2_(µ- SCH_2_CHCH_2_S)(CO)_4_(CN)_2_]. This alternative mimic was synthesised according to previous protocols with minor modifications and verified by FTIR spectroscopy^78–80^. The dense cell suspension was centrifuged, and the cell pellet was washed with 1 mL Tris-HCl buffer (100 mM Tris, 150 mM NaCl, pH 8.0) three times under anaerobic conditions. The cells were then resuspended with 200 µL Tris buffer pH 8.0 and transferred into EPR tubes. The tubes were capped and promptly frozen in liquid N_2_. Measurements were performed on a Bruker ELEXYS E500 spectrometer using an ER049X SuperX microwave bridge in a Bruker SHQ0601 cavity equipped with an Oxford Instruments continuous flow cryostat and using an ITC 503 temperature controller (Oxford Instruments).

### Metagenomic analyses across health status

To assess the distribution of hydrogenases across health status, we used a previously curated and quality controlled dataset containing 1661 metagenomes from 33 studies^55^. The dataset encompassed 871 healthy and 790 diseased individuals, including 11 chronic disease phenotypes. Quality control was performed with TrimGalore v.0.6.6^111^ using a threshold of 80 bp for read length and minimum Phred score of 25. Host sequence reads were removed by mapping the sequence reads to the human genome with bowtie v.2.3.552^112^. To minimize the impact of sequence depth, samples were rarefied to 15M reads with seqtk v.1.3^113^, as previously described^55^. Forward reads were mapped to a dataset of hydrogenase and ribosomal RNA sequences with DIAMOND v2.0^93^. as described above. Alignments were filtered to a minimum length of 26 amino acids, subject to identity threshold filtering, and normalized to gene copy per organism as described above. The largest case-control IBD-related study within this dataset^88^ was selected to investigate the distribution of hydrogenase subgroups between healthy and disease-associated microbiomes, which included 46 patients with Crohn’s disease and 38 healthy controls. Statistical significance was assessed with Wilcoxon tests, using the Holm–Bonferroni method to account for multiple comparisons across disease states.

## Acknowledgements

This study was supported by an NHMRC EL2 Fellowship (APP1178715; to C.G.) and an Australian Government Research Training Scholarship (to C. W.). R.L. and V.R.M are supported by ARC DECRA Fellowships (DE230100542 and DE220100965). D.L. is supported by Australian Research Council Laureate Fellowship (FL210100258).

## Footnotes

### Author contributions

C.G., C.W., S.C.F., D.L., P.W., H.R.G., and J.R. conceptualised this study. C.G., S.C.F., D.L., C.W., R.L., T.D.W., and G.B. supervised students. C.G., C.W., S.C.F., G.B., R.L., and T.D.W. designed experiments. C.W. conducted most experiments and analysed most data. Specific authors contributed to metagenomic screening (C.W., C.G., R.L.), genomic screening (C.W., S.C.F., E.L.G.), biopsy metagenome collection (G.D.A., E.M.G., S.C.F., R.B.Y.), culture-based gas assays (C.W., N.Q.D., N.B., J.S, E.L.G., S.C.F.,C.G.), transcriptomics (C.W., D.J.K., J.A.G., S.C.F., T.D.W., C.G.), biochemical characterisation (P.C., G.B., P.H., K.W., C.G.), structural modelling (J.L., C.G., D.J.K.), and analyses of disease states (V.R.M., C.G., C.W., S.C.F., R.B.Y.). C.W., C.G., T.D.W., and R.L. wrote the paper with input from all authors.

### Data availability

The new metagenomes, genomes, and transcriptomes analysed in this study will be available prior to publication.

## Supplementary information

**Supplementary Note 1. EPR characteristics of the *Bacteroides* [FeFe]-hydrogenases.** The [2Fe]^pdt^ cofactor mimic lacks the nitrogen bridgehead of [2Fe]^adt^, hampering catalysis and promoting the build-up of the EPR-active H-cluster resting state, H_ox_. The [2Fe]^pdt^-treated whole cells expressing the *B. fragilis* group B [FeFe]-hydrogenase showed a partial rhombic signal for H_ox_-like state, with the first two g-values in good agreement with previously reported [FeFe]- hydrogenases, typically with values above *g* = 2^·^ (see **Table S6**). The third *g*-value is theoretically positioned at around 2.010-2.009, but it cannot be distinctly resolved due to strong overlap with BL21(DE3) cell background signals. Whole-cell samples of the *B. thetaiotaomicron* group B [FeFe]-hydrogenase, the second most active gut hydrogenase in the activity screening, did not exhibit clear H-cluster signals when incubated with [2Fe]^pdt^ and exhibited a spectrum equivalent to that of BL21(DE3) cells not expressing any [FeFe]- hydrogenase (data not shown). The absence of any discernible EPR signal attributable to the H-cluster in these samples is potentially due to the formation of thermodynamically favourable H-cluster states that are EPR-silent, but is more likely due to the low solubility and thereby low concentration of holo-enzyme in the whole-cell mixture.

---

**Figure S1.**
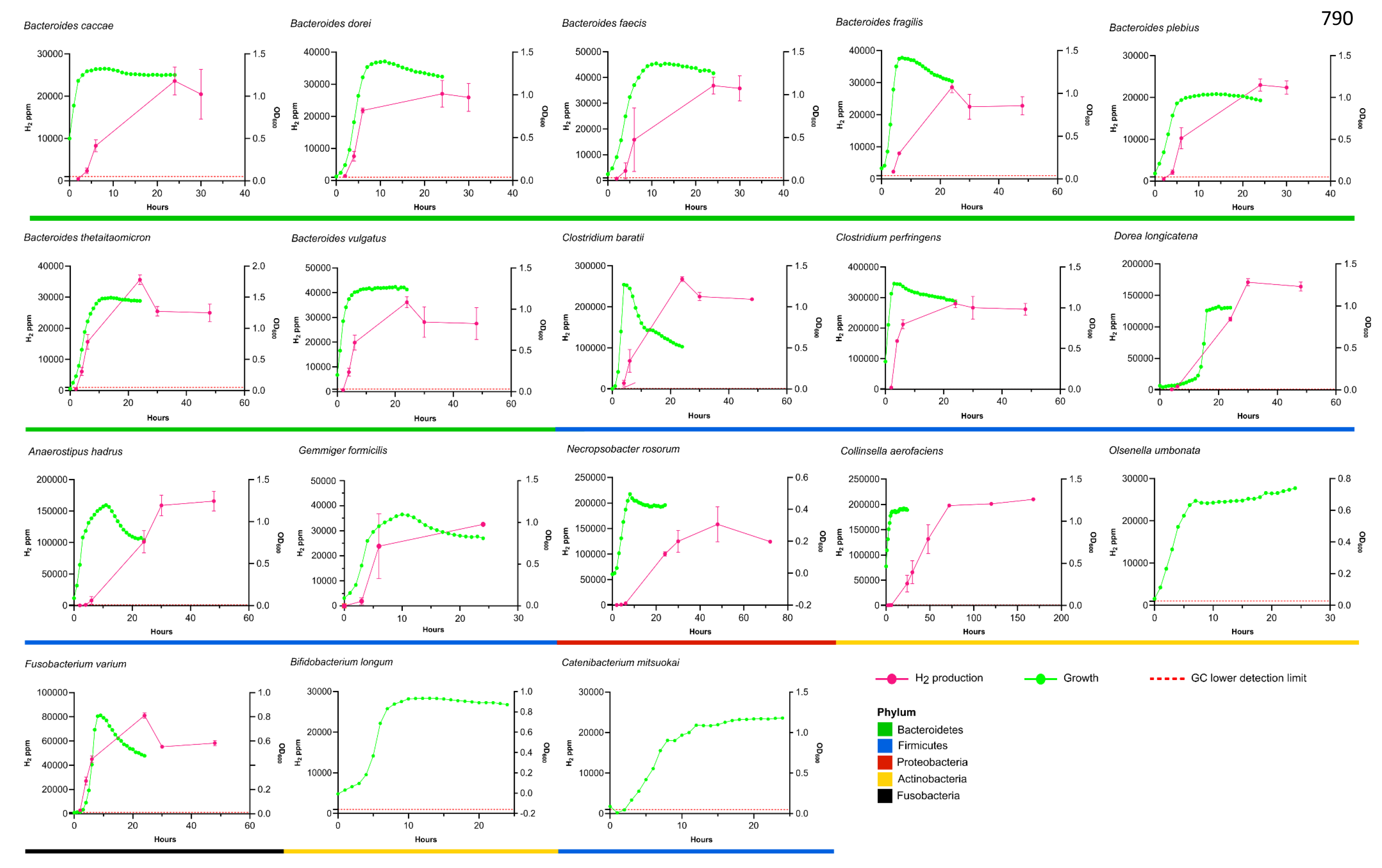
**Comparison of the growth and H2 production of the 18 human gut isolates**

**Figure S2.**
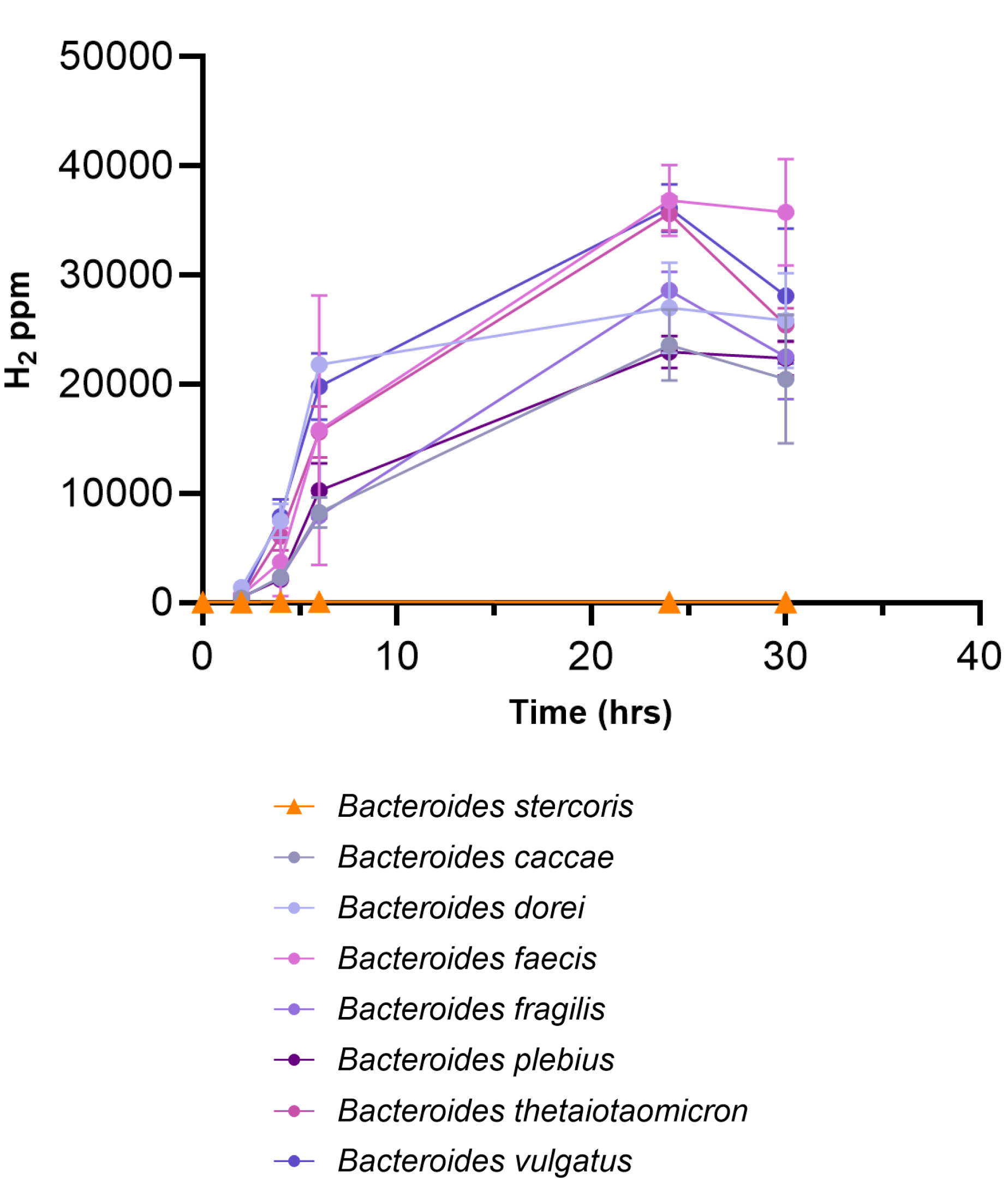
Comparison of H_2_ production activities in *Bacteroides* strains containing and lacking group B [FeFe]-hydrogenases. The *B. stercoris* strain lacks group B [FeFe]- hydrogenases, whereas the other strains encode and express them.

**Figure S3.**
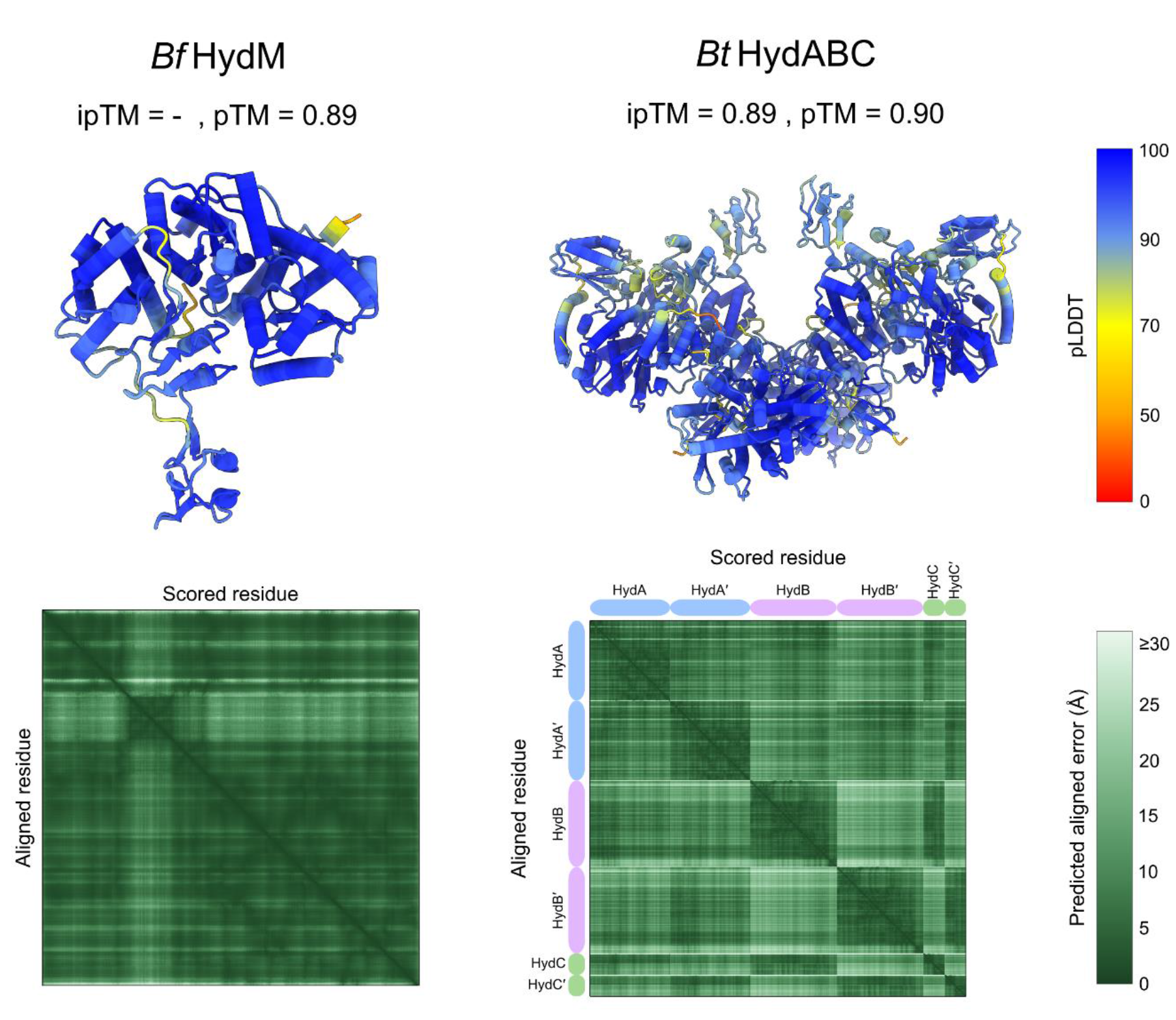
AlphaFold2 predicted protein structures. AlphaFold2 confidence scores for the two protein models made in this study. The top ranked model is shown and coloured according to their predicted local distance difference test (pLDDT) score, and their corresponding predicted aligned error (PAE) plots shown below. Portions of the *Bt*HydABC PAE plot axis are labelled according to their subunit identities. PAE plots were generated using PAE Viewer web server^114^. ipTM: interface predicted template modelling score. pTM: predicted template modelling score.

**Figure S4.**
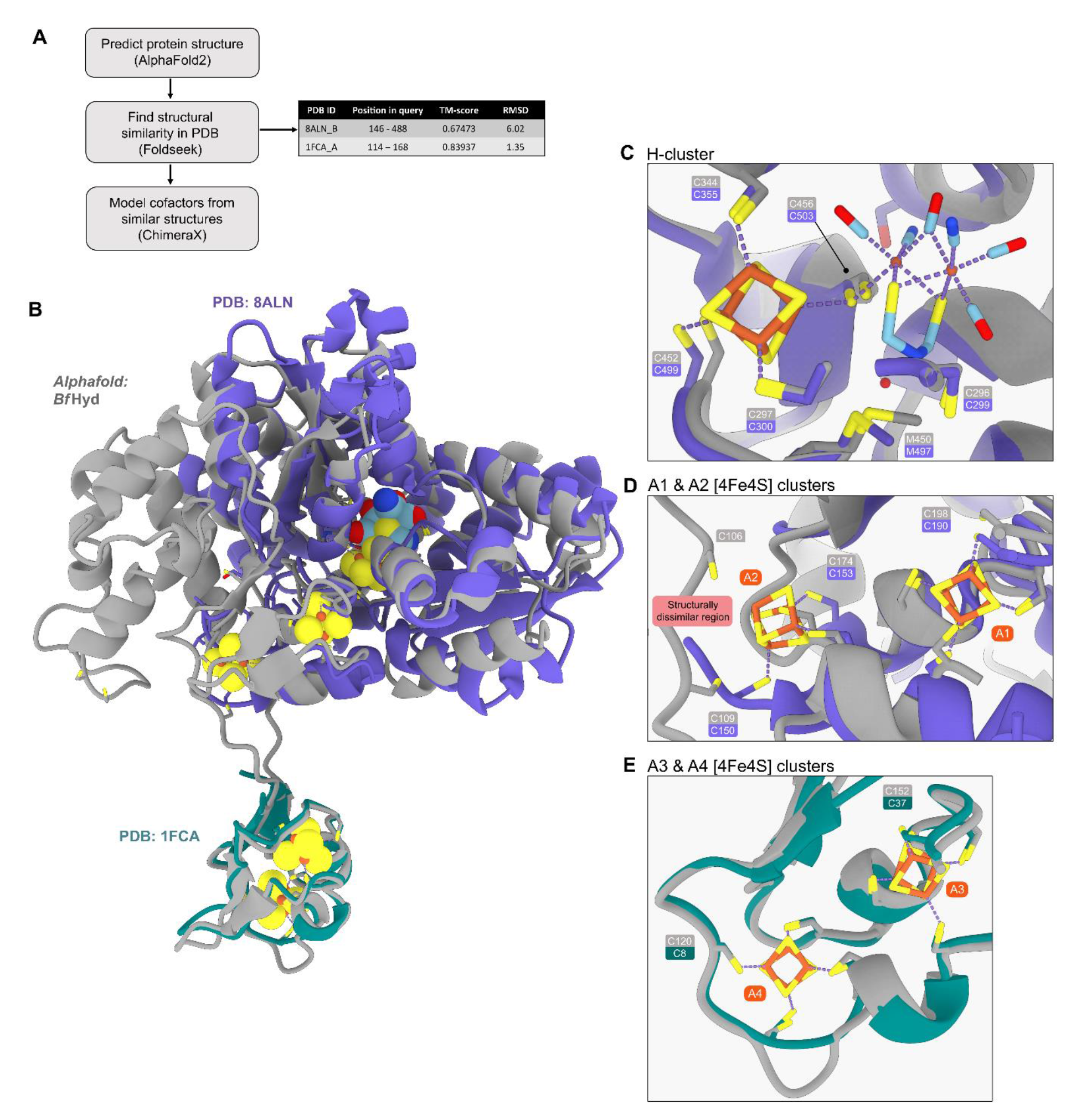
Process for modelling putative cofactors into the predicted structure of the. ***B. fragilis* group B [FeFe]-hydrogenase predicted structure. (a)** Overview of strategy used for modelling cofactors into apo *Bf*HydM. Experimental structures chosen from FoldSeek output summarised on the right with their corresponding positional overlap and structural overlap (TM-score and RMSD). **(b)** Superposition of apo *Bf*HydM against overlapping portions of experimental structures chosen from FoldSeek. The PDBs were chosen to act as a template for cofactor modelling based on their (i) presence of experimentally observed cofactors and (ii) structural similarity to portions of apo *Bf*HydM, especially at conserved cysteine residues. **(c)** the putative H-cluster pocket in *Bf*HydM is highly similar to the structural architecture of that seen in the group A1 [FeFe]-hydrogenase of *Clostridium pasteurianum* (*Cp*I; PDB: 8ALN^108^). Conserved H-cluster binding residues between the two structures overlay near identically, allowing for minimal manual repositioning of the H-cluster into *Bf*HydM. **(d)** Putative iron-sulfur cluster pockets (A1 and A2) in *Bf*HydM share some structural similarity to those seen in group A1 [FeFe]-hydrogenase from *Cp*I, but with larger divergence compared to the aforementioned H-cluster. The iron-sulfur clusters required manual repositioning to coordinate with the cysteines and bond lengths were checked as being biochemically reasonable in the UCSF ChimeraX software. **(e)** Putative iron-sulfur cluster pockets (A3 and A4) in *Bf*HydM share near identical structural similarity to those seen in *Cp*I. Iron-sulfur clusters were transposed into *Bf*HydM without any manual repositioning from their relative positions observed in the *Cp*I structure.

**Figure S5.**
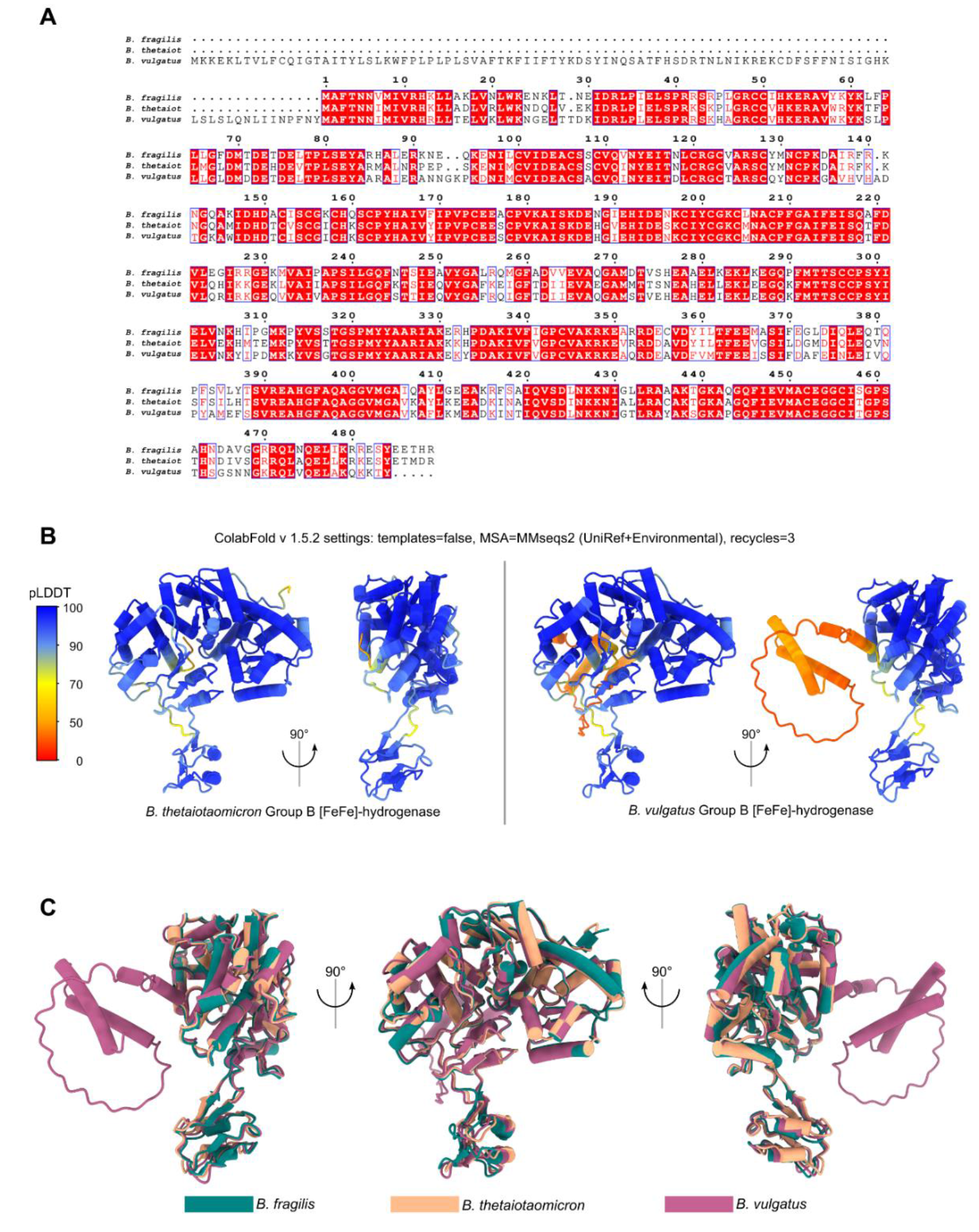
AlphaFold2 structural predictions of group B [FeFe]-hydrogenases from the *Bacteroides*. (a) Sequence conservation of Group B [FeFe]-hydrogenase homologs between *Bacteroides fragilis*, *Bacteroides thetaiotaomicron*, and *Bacteroides vulgatus*. Multiple sequence alignment was performed with ClustalO v1.2.4 and visualised with the ESPript 3.0 web server^115,116^. **(b)** Top-ranked AlphaFold2 models of the *B. thetaiotaomicron* and *B. vulgatus* group B [FeFe]-hydrogenase, coloured by pLDDT. The low pLDDT scoring N-term portion of *Bv*Hyd is possibly an intrinsically disordered domain. **(c)** Superposition of all three *Bacteroides* Group B [FeFe]-hydrogenases showing overall structural conservation, except for the N-terminal portion of *Bv*HydM which is predicted to be largely disordered.

**Figure S6.**
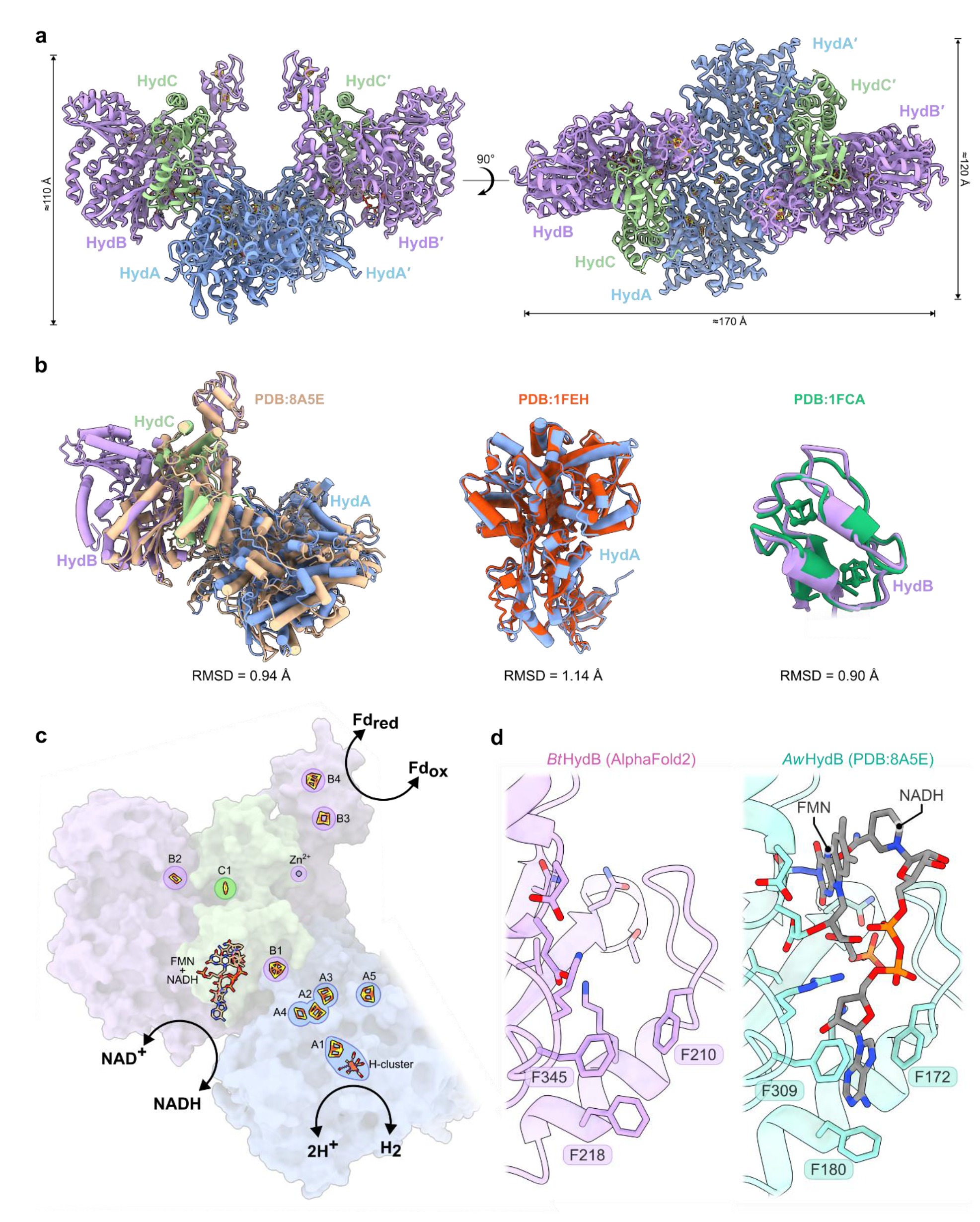
Predicted structure of the group A3 [FeFe]-hydrogenase from *Bacteroides thetaiotaomicron*. (a) Top and side view of the AlphaFold2 predicted structure. **(b)** Superposition of HydABC components with structures of *Acetobacterium woodii* HydABC (PDB ID: 8A5E^64^), *Clostridium pasteurianum* [FeFe]-hydrogenase (PDB ID: 1FEH^40^), and *Clostridium acidurici* ferredoxin (PDB ID: 1FCA^109^). The structural similarity between these proteins and *Bt*HydABC allowed for homology modelling of cofactors. **(c)** Putative cofactors and enzymatic reactions of *Bt*HydABC. Cofactors are positioned based on homology models and labelled according to their subunit identity. **(d)** Structure of the putative FMN and NADH binding site in the AlphaFold structure (left) compared to the same site in the experimental structure of *A. woodii* HydB (right). A trio of phenylalanine residues which form a π-stacking “clamp” around the adenine moiety of NADH is conserved in both structures. RMSD: root- mean-square-deviation. Å: ångström. FMN: flavin mononucleotide. NADH: nicotinamide adenine dinucleotide (reduced). NAD^+^: nicotinamide adenine dinucleotide (oxidized). Fd_red_: reduced ferredoxin. Fd_ox_: oxidized ferredoxin.

**Figure S7.**
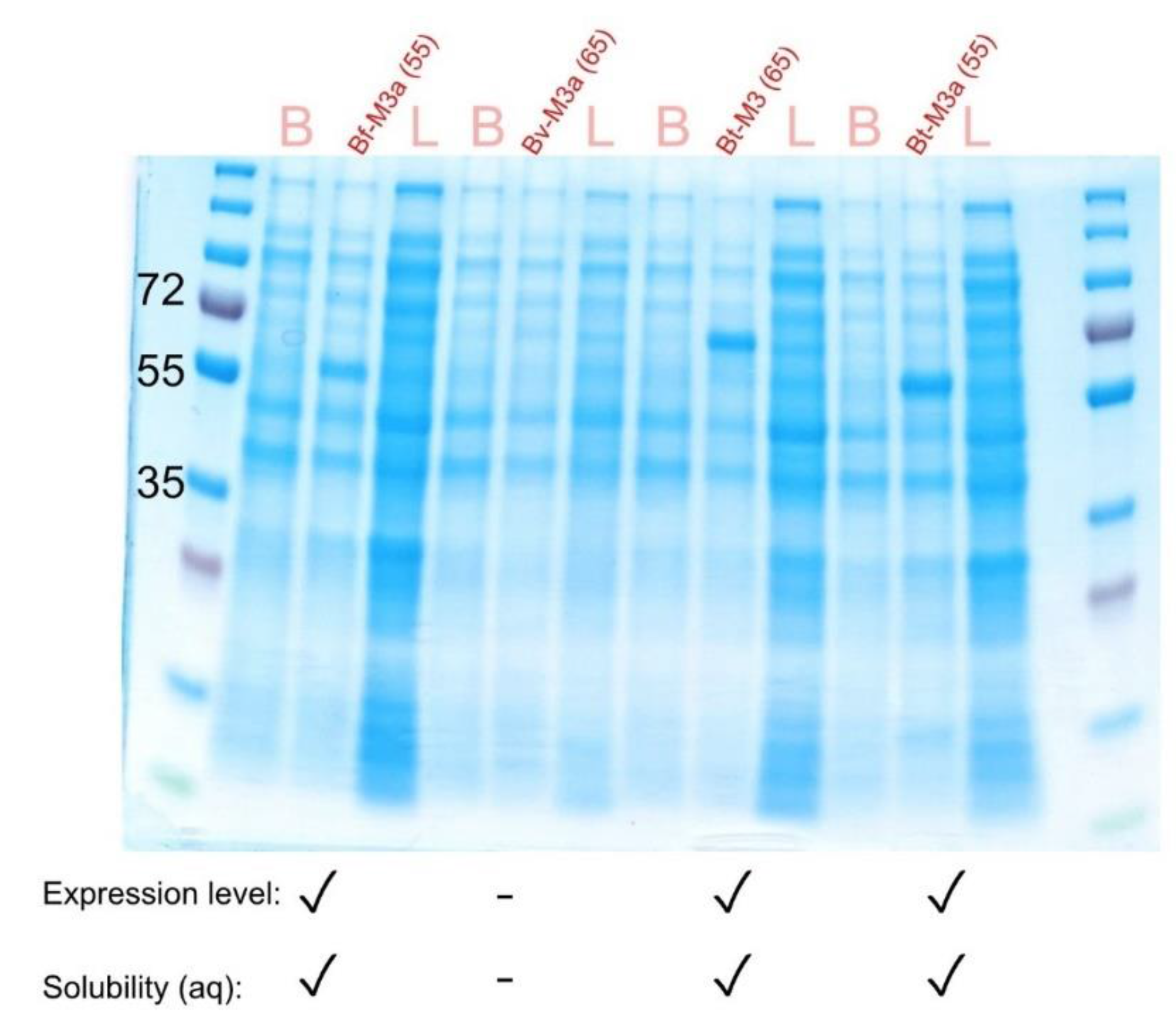
SDS-PAGE visualising the molecular weights of the heterologously expressed [FeFe]-hydrogenases from *Bacteroides*. Expression constructs with verified sequences were used to transform chemically competent *E. coli* BL21(DE3). Protein bands are shown from before induction with IPTG (B), after induction (Name-Subclass and with the expected kDa size in parenthesis), and lysate or supernatant after cell lysis and centrifugation (L). The bands in each after-induction lane corresponded well with the expected molecular weights in kDa. Three gut-associated [FeFe]-hydrogenases (**Bf-M3a**, **Bt-M3,** and **Bt-M3a**) exhibited high levels of expression and low to moderate solubility while **Bv-M3a** had poor expression and solubility levels.

**Figure S8.**
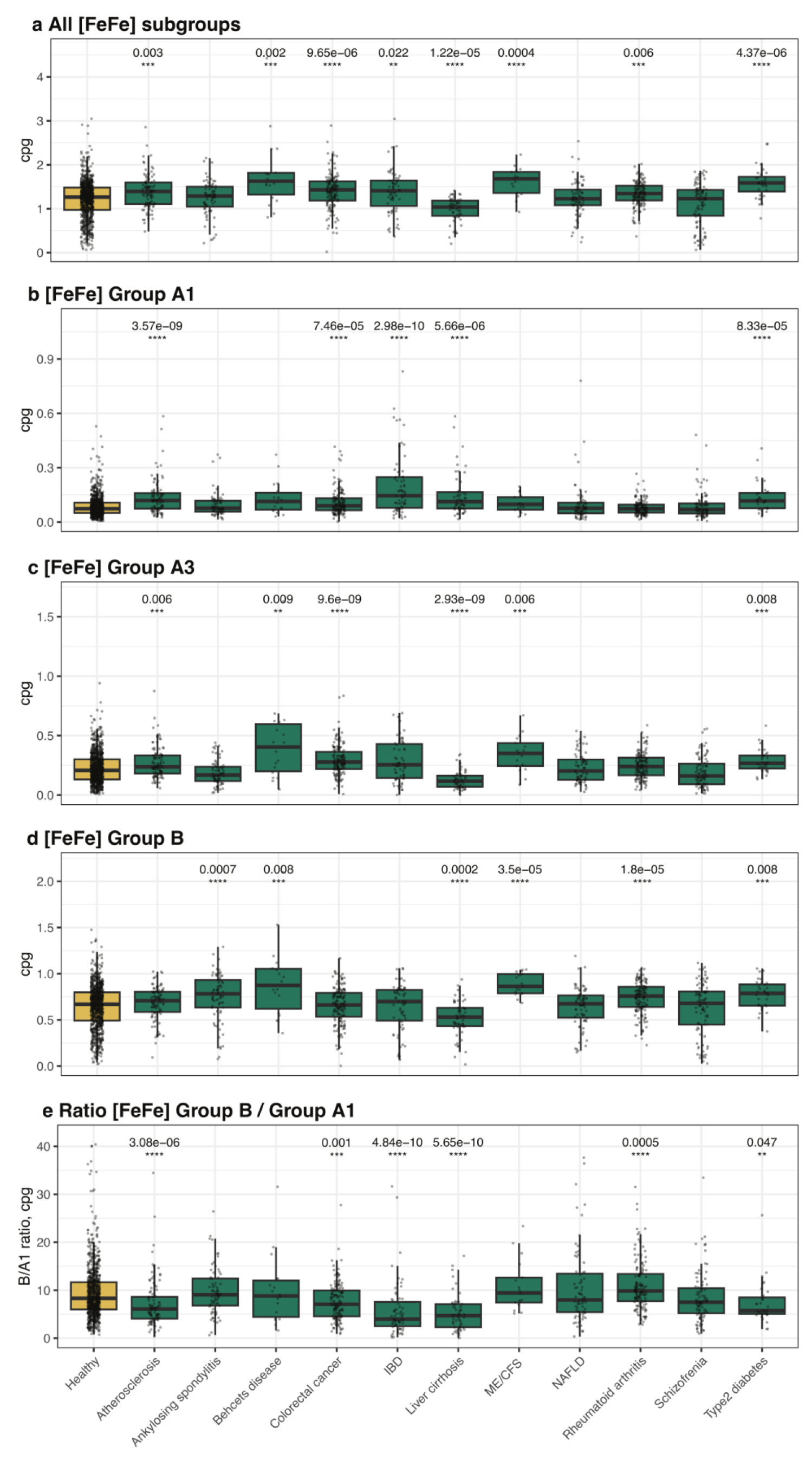
Distribution of key [FeFe] hydrogenase subgroups across diseases. Statistical significance was assessed with Wilcoxon tests, using the Holm–Bonferroni method to account for multiple comparisons. IBD = inflammatory bowel disease. cpg = counts per genome. ME/CSF = myalgic encephalomyelitis / chronic fatigue syndrome.

**Figure S9.**
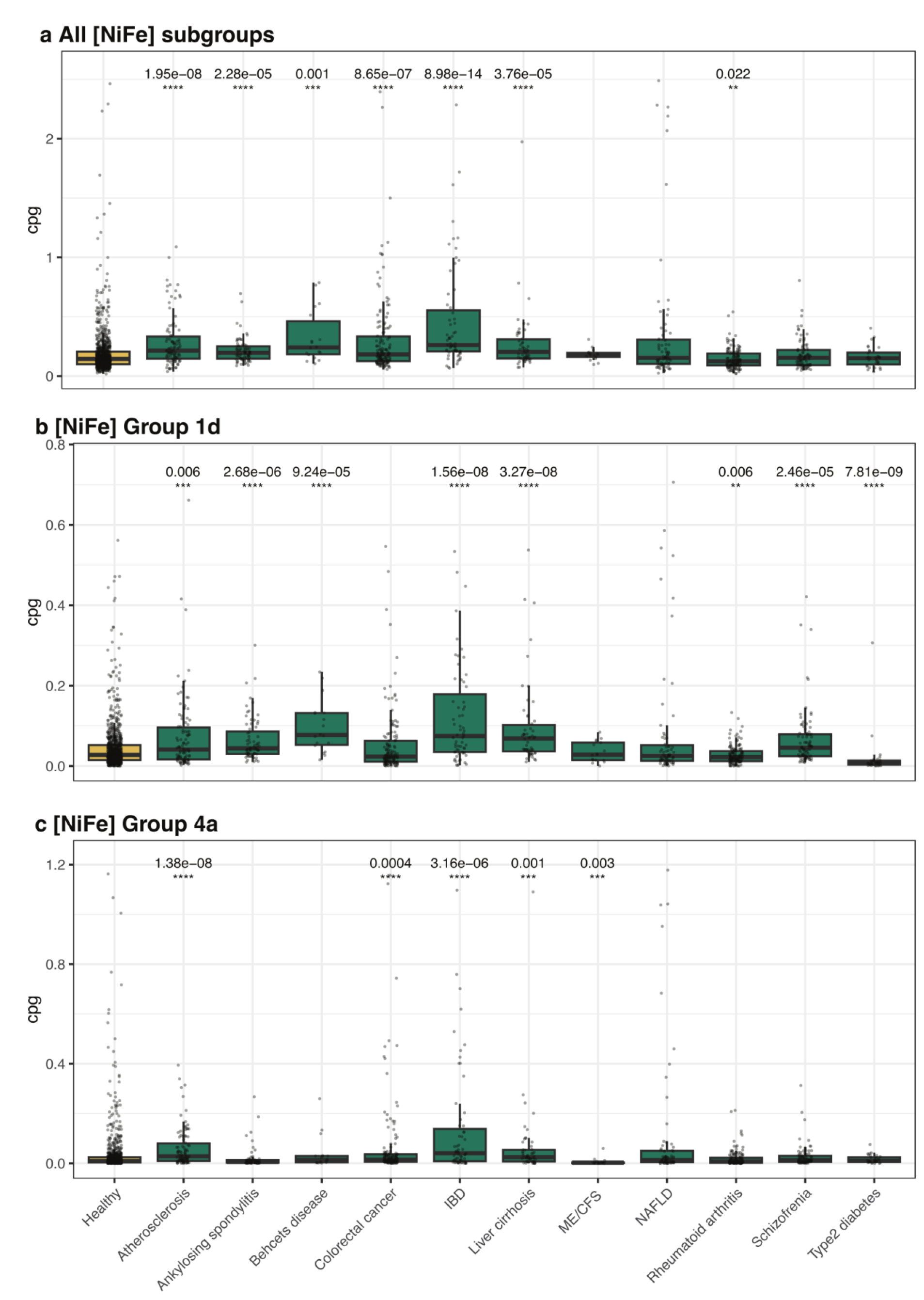
Distribution of key [NiFe] hydrogenase subgroups across diseases. Statistical significance was assessed with Wilcoxon tests, using the Holm–Bonferroni method to account for multiple comparisons. IBD = inflammatory bowel disease. cpg = counts per genome. ME/CSF = myalgic encephalomyelitis / chronic fatigue syndrome.

**Table S1 (xlsx)**. Abundance, expression, and origin of hydrogenases and associated genes in stool metagenomes, stool metatranscriptomes, and biopsy metagenomes.

**Table S2 (xlsx)**. Taxonomy and metabolic capabilities of the 812 sequenced human gut isolates from the Australian Microbiome Culture Collection (AusMiCC).

**Table S3.** Taxonomy and hydrogenase content of the 19 gut isolates from Australian Microbiome Culture Collection (AusMiCC) used for growth, gas, and transcriptome analyses.

| Species | Strain | Hydrogenase subgroup |
| --- | --- | --- |
| *Clostridium perfringens* | CC01445 | Group B [FeFe]-hydrogenase<br>Group B [FeFe]-hydrogenase<br>Group A1 [FeFe]-hydrogenase<br>Group A1 [FeFe]-hydrogenase |
| *Clostridium baratii* | CC01452 | Group B [FeFe]-hydrogenase<br>Group A1 [FeFe]-hydrogenase |
| *Fusobacterium varium* | CC01421 | Group A1 [FeFe]-hydrogenase<br>Group A1 [FeFe]-hydrogenase<br>Group A3 [FeFe]-hydrogenase |
| *Necropsobacter rosorum* | CC01403 | Group 1c [NiFe]-hydrogenase<br>Group 4a [NiFe]-hydrogenase |
| *Bacteroides caccae* | CC01389 | Group B [FeFe]-hydrogenase<br>Group A3 [FeFe]-hydrogenase |
| *Bacteroides thetaiotaomicron* | CC00765 | Group B [FeFe]-hydrogenase<br>Group A3 [FeFe]-hydrogenase |
| *Bacteroides dorei* | CC01440 | Group B [FeFe]-hydrogenase |
| *Bacteroides faecis* | CC01412 | Group B [FeFe]-hydrogenase<br>Group A3 [FeFe]-hydrogenase |
| *Bacteroides vulgatus* | CC01422 | Group B [FeFe]-hydrogenase |
| *Bacteroides fragilis* | CC01400 | Group B [FeFe]-hydrogenase |
| *Bacteroides plebius* | CC01397 | Group B [FeFe]-hydrogenase |
| *Gemmiger formicilis* | CC00311 | Group B [FeFe]-hydrogenase<br>Group A1 [FeFe]-hydrogenase<br>Group A2 [FeFe]-hydrogenase |
| *Dorea longicatena* | CC00515 | Group B [FeFe]-hydrogenase<br>Group A2 [FeFe]-hydrogenase |
| *Anaerostipes hadrus* | CC00501 | Group B [FeFe]-hydrogenase<br>Group B [FeFe]-hydrogenase<br>Group A2 [FeFe]-hydrogenase |
| *Olsenella umbonata* | CC00540 | Group B [FeFe]-hydrogenase<br>Group A2 [FeFe]-hydrogenase |
| *Collinsella aerofaciens* | CC00529 | Group 4a [NiFe]-hydrogenase<br>Group 4e [NiFe]-hydrogenase<br>Group A2 [FeFe]-hydrogenase |
| *Bifidobacterium longum* | CC00565 | None |
| *Catenibacterium mitsuokai* | CC00599 | None |
| *Bacteroides stercoris* | CC00654 | None |

**Table S4 (xlsx)**. Annotated transcriptomes of the 18 gut isolates.

**Table S5.** Features and activities of the [FeFe]-hydrogenases heterologously expressed and semisynthetically matured in *E. coli*.

| Group | Subclass | Species | [FeS] Architecture | Size (kDa) | % Activity* |
| --- | --- | --- | --- | --- | --- |
| B | M3a | *Bacteroides fragilis* (Bf) | 3 x [4Fe-4S], 1 x [2Fe-2S] | 55 | 34 ± 6 |
| B | M3a | *Bacteroides vulgatus* (Bv) | 3 x [4Fe-4S], 1 x [2Fe-2S] | 65 | 1.9 ± 0.3 |
| B | M3a | *Bacteroides thetaiotaomicron* (Bt) | 3 x [4Fe-4S], 1 x [2Fe-2S] | 55 | 7.6 ± 1.3 |
| A3 | M3 | *Bacteroides thetaiotaomicron* (Bt) | 3 x [4Fe-4S], 1 x [2Fe-2S] | 65 | 0.22 ± 0.04 |

\* with respect to the H<sub>2</sub> evolution rates of the *Chlamydomonas reinhardtii* group A1 [FeFe]-hydrogenase (CrHydA1).

**Table S6.** Previously reported *g*-values for [FeFe]-hydrogenases matured with [2Fe]^pdt^, i.e. H_ox_- [2Fe]^pdt^ state.

| Group | Subclass | Species | *g*<sub>1</sub> | *g*<sub>2</sub> | *g*<sub>3</sub> |
| --- | --- | --- | --- | --- | --- |
| A1 | M1 | *Chlamydomonas reinhardtii* (CrHydA1)<sup>14</sup> | 2.094 | 2.039 | 1.998 |
| A1 | M3 | *Clostridium pasteurianum* (Cpl)<sup>15</sup> | 2.092 | 2.039 | 2.000 |
| A1 | M2 | *Solobacterium moorei* (SmHydA)<sup>4</sup> | 2.100 | 2.040 | 2.010 |
| A1 | M2 | *Desulfovibrio desulfuricans* (DdH)<sup>16</sup> | 2.095 | 2.041 | 1.998 |
| B | M3a | ***Bacteroides fragilis* (BfHydM, this study)** | **2.101** | **2.053** | **Not detected** |
| C | M2f | *Thermotoga maritima* (TmHydS)<sup>17</sup> | 2.108 | 2.043 | 2.000 |
| D | M2e | *Thermoanaerobacter mathranii* (TamHydS)<sup>4</sup> | 2.106 | 2.051 | 2.010 |

**Table S7 (xlsx)**. Hydrogenase levels in the stool metagenomes of healthy individuals compared to those with chronic disease states.

